# CPI-Pred: A deep learning framework for predicting functional parameters of compound-protein interactions

**DOI:** 10.1101/2025.01.16.633372

**Authors:** Zhiqing Xu, Rana Ahmed Barghout, Jinghao Wu, Dhruv Garg, Yun S. Song, Radhakrishnan Mahadevan

## Abstract

Recent advancements in deep learning have enabled functional annotation of genome sequences, facilitating the discovery of new enzymes and metabolites. However, accurately predicting compound-protein interactions (CPI) from sequences remains challenging due to the complexity of these interactions and the sparsity and heterogeneity of available data, which constrain the generalization of patterns across their solution space. In this work, we introduce CPI-Pred, a versatile deep learning model designed to predict compound-protein interaction function. CPI-Pred integrates compound representations derived from a novel message-passing neural network and enzyme representations generated by state-of-the-art protein language models, leveraging innovative sequence pooling and cross-attention mechanisms. To train and evaluate CPI-Pred, we compiled the largest dataset of enzyme kinetic parameters to date, encompassing four key metrics: the Michaelis-Menten constant (*K*_M_), enzyme turnover number (*k*_cat_), catalytic efficiency (*k*_cat_*/K*_M_), and inhibition constant (*K*_I_).These kinetic parameters are critical for elucidating enzyme function in metabolic contexts and understanding their regulation by compounds within biological networks. We demonstrate that CPI-Pred can predict diverse types of CPI using only the amino acid sequence of enzymes and structural representations of compounds, outperforming state-of-the-art models on unseen compounds and structurally dissimilar enzymes. Over workflow provides a valuable tool for tackling a range of metabolic engineering challenges, including the designing of novel enzyme sequences and compounds, such as enzyme inhibitors. Additionally, the datasets curated in this study offer a valuable resource for the scientific community, serving as a benchmark for machine learning models focused on enzyme activity and promiscuity prediction.

## Introduction

Deep learning has exhibited exceptional potential in recent years to advance systems biology and metabolic engineering. The rapid growth of bioinformatics databases has not only enhanced our understanding of genome organization and protein function but also catalyzed the development of computational tools for engineering biology, including protein design. Advancements in high-performance computing (HPC) and the increasing availability of computational resources have enabled the integration of diverse algorithms into systematic protein design pipelines, achieving progressively better performance^1,2^.

Recent deep learning predictive models, such as AlphaFold^1^, Chai-1^3^, and OpenFold^2^, have demonstrated remarkable accuracy in predicting three-dimensional (3D) protein structures from amino acid sequences. These models employ state-of-the-art algorithms to identify the spatial arrangement of atoms within proteins with accuracy comparable to experimental methods. Their unprecedented effectiveness highlights the potential of deep learning techniques to deepen our understanding of protein properties and functions. Similarly, self-supervised language models, including TAPE^4^, ProtBERT^5^, ESM^6^, Ankh^7^, and CARP^8^, leverage entire protein databases to generate embeddings that capture contextual and structural protein information. These embeddings have been shown to enhance sequence-to-function predictions, improving tasks such as protein contact map prediction, fluorescence, and stability^4^.

Despite the significant progress made in various supervised (and semi-supervised) learning tasks for protein biology, compound-protein interaction (CPI) prediction remains an active area of research for its diverse range of applications including drug discovery, systems biology, biochemical production, and environmental toxicology, etc. Therefore, it is imperative to address the underlying shortcomings of existing CPI prediction frameworks to improve accuracy and robustness. The difficulty of CPI prediction arises from several reasons, including, (1) the complexity of interactions between compounds and proteins (which involve multiple binding sites, different binding specificity and affinity, etc.) and (2) the sparsity and heterogeneity of data sources (insufficient data to fully capture the range of features), which makes it difficult to generalize patterns across the solution space ^9,10^.

In the realm of CPI prediction, several models have made significant strides. The CLEAN model, a contrastive learning model, has been particularly effective in predicting high-quality functional properties of proteins^11^. It uses enzyme commission (EC) numbers to provide accurate, reliable, and sensitive annotations, outperforming existing computational tools. Building on CPI prediction advancements specific to enzyme annotation, the ESP model emerges as a robust deep learning framework, achieving over 91% accuracy in predicting enzyme-substrate pairs across diverse enzymes and metabolites ^12^.

In addition to these advancements in predictive modeling, understanding enzyme behaviour through kinetic parameters is crucial for comprehensive protein characterization. Kinetic parameters are mathematical constants that describe the rate and efficiency of enzyme-catalyzed reactions, commonly determined experimentally using a variety of techniques, such as enzyme assays, spectroscopy, and chromatography. The most commonly studied kinetic parameter is the Michaelis-Menten constant (*K*_M_), a measure of the substrate concentration required to achieve half of the maximum reaction rate. Other important kinetic parameters include the enzyme turnover number (*k*_cat_), which is a measure of the number of substrate molecules converted to product per unit time, and the catalytic efficiency (*k*_cat_*/K*_M_), which is a measure of the enzyme’s ability to convert substrate to product per unit of substrate concentration. The *K*_I_ value, or inhibition constant, is a measure of the binding affinity between an enzyme and an inhibitor. It is commonly used to evaluate the potency of enzyme inhibitors and to compare the relative affinity of different inhibitors.

Building on the importance of kinetic parameters, another noteworthy model is the one developed by Kroll et al^13^. This model uses a deep learning framework to predict the Michaelis constant (*K*_M_) for wild-type enzyme-substrate combinations. The model was trained on a dataset with 11,675 entries of wild-type enzymes, demonstrating the potential of deep learning to predict enzyme kinetics . These models, each with their unique approaches and strengths, contribute to the ongoing advancements in CPI predictions, although there is still significant work to be done. Complementing the deep learning framework by Kroll et al., the DLkcat model developed by Li et al. represents a significant advancement in enzyme kinetics prediction ^14^. This model facilitates high-throughput *k*_cat_ prediction for metabolic enzymes from any organism, utilizing substrates structures and protein sequences.

Although many of these CPI prediction models obtain qualitative function predictions while utilizing GNNs and transformer-based architectures, there is no deep learning framework that can predict all compound-protein parameters in a unified manner. Furthermore, CPI prediction models have achieved some success in predicting enzyme (sequence) variant activity on new substrates^9,15^, but investigations have found that deep learning of *k*_cat_ values or catalytic ability of proteins on compounds is sometimes outperformed by simple machine learning baseline models, such as the K-nearest neighbors (KNN) model ^10^. This is due, in part, to downstream predictive models receiving overly compressed compound representations and low-grade sequence embeddings method ^14^. The KNN algorithm relies on identifying the most similar data points in the training set and uses this information to make predictions for new data points. KNN models outperforming deep learning-based approaches suggests that low-grade compound and sequence representations are not able to capture the similarities between data points. Here, we present CPI-Pred, a deep learning model that extends a previous enzyme activity prediction pipeline ^9^ with (1) updated protein language models and improved sequence embedding pooling algorithm, (2) a learned representation of compounds via a message-passing neural network with customized atom-level features, (3) a cross-attention block for better interaction prediction to replace previously used concatenation of compound and sequence representations, and (4) an ensemble of multiple prediction pipelines to further increase the robustness and accuracy of the model. We demonstrate the broad utility of this model architecture by showing that it achieves good performance with different protein embeddings and outperforms other existing models (i.e. DLkcat^14^, HAC-Net^16^) and a non-neural network baseline.

Apart from an improved deep learning-based workflow, the availability of diverse and high-quality data is also essential for developing accurate and robust CPI prediction models. While there are several high-throughput experimental techniques available to study compound protein interactions, they are often costly and time-consuming to carry out^17^. Thus, collecting large, well-labeled datasets requires extensive experimental workflows and is, therefore, a major bottleneck. While data augmentation approaches have been used to expand existing datasets^18^, the synthetic dataset generated from a original dataset with limited size may not accurately represent the true distribution of the data^19^. Such limitations may hinder the ability of the augmented dataset to effectively capture the full range of variability and complexity of the underlying data.

Accurate determination of kinetic parameters is essential for understanding enzyme function, designing inhibitors or activators of enzyme activity, and developing quantitative models of metabolic pathways and cellular processes for synthetic biology applications including enzyme and metabolic pathway engineering. These four kinetic parameters are functional descriptors of compound-protein interaction, and in this work we use them to evaluate the model performance through a rigorous task assessment pipeline.

The kinetic parameters explored in this study were extracted from the Braunschweig Enzyme Database (BRENDA)^20^. One challenge with these large datasets is that approximately 60% of the data from BRENDA lacks UniProt IDs, and therefore introduces ambiguity in relation to the protein sequences. In order to utilize these instances with unannotated UniProt IDs^21^, we leverage the KinMod database^22^, which adopts a ”pangenomic” approach and provides detailed information on the relationships between sequences, organisms, and Enzyme Commission (EC) numbers. Using this database, we assign sequences to data points that lack UniProt IDs based on the associated organism and EC number. Since the extended datasets (hereinafter referred to as pangenomic datasets) obtained using this approach are subject to some uncertainty, we perform an extended screening process to ensure data quality. To validate the pangenomic kinetic parameters datasets, we demonstrate the performance improvements achieved by using these datasets to train the model. Furthermore, we evaluate the model’s sequence-to-function prediction ability on two experimental datasets^17,23^ as well as functions beyond kinetic parameter prediction, such as binding affinity using a highquality dataset (PDBbind)^24,25^. The PafA dataset^17^ provides a case study for the ability of CPI-Pred to learn from enzyme variant data obtained from a high-through put mutational scanning technique highlighting the value for using CPI-Pred for enzyme engineering whereas the Sarkisyan et al.^23^ dataset is a classic benchmark to predict GFP fluorescence, that is used as a test case in many protein property prediction pipelines. All the ten CPI datasets developed in this study represent diverse case studies including prediction of enzyme kinetics, binding affinity, enzyme variant activity and protein property and are summarized in Supplementary Table S1 and serve as a valuable bench-marking datasets for future algorithms similar to the ImageNet database^26^.

## Methods

In this study, we first establish a comprehensive database of kinetic parameters. Following this, we develop a deep learning workflow that integrates novel methods for representing compounds and protein sequences, utilizing self-attention and cross-attention mechanisms to combine these representations. We evaluate the effectiveness of this pipeline across various tasks, including prediction for unseen enzymes and compounds. Additionally, we implement an error predictor to provide not only the predicted values of kinetic parameters, but also the associated error levels, enabling users to assess and utilize the predictions more effectively.

In this section, we outline the approach employed to predict compound-protein interactions, covering various aspects including data processing, sequence and compound encoding, and interaction modeling. We discuss the pangenomic approach to obtain large kinetic parameter datasets as well as the usage of the KinMod database in the data processing pipeline, followed by the validation of the screened pangenomic dataset. We elaborate on the architecture of the CPI prediction pipeline as shown in Figure 1A. Additionally, we describe different sequence embedding pooling methods for representing proteins and combine this with a compound encoding based on a message passing neural network (MPNN) with a novel atom-level featurization. Finally, we introduce a cross-attention approach for learning compound-protein interactions.

**Figure 1:**
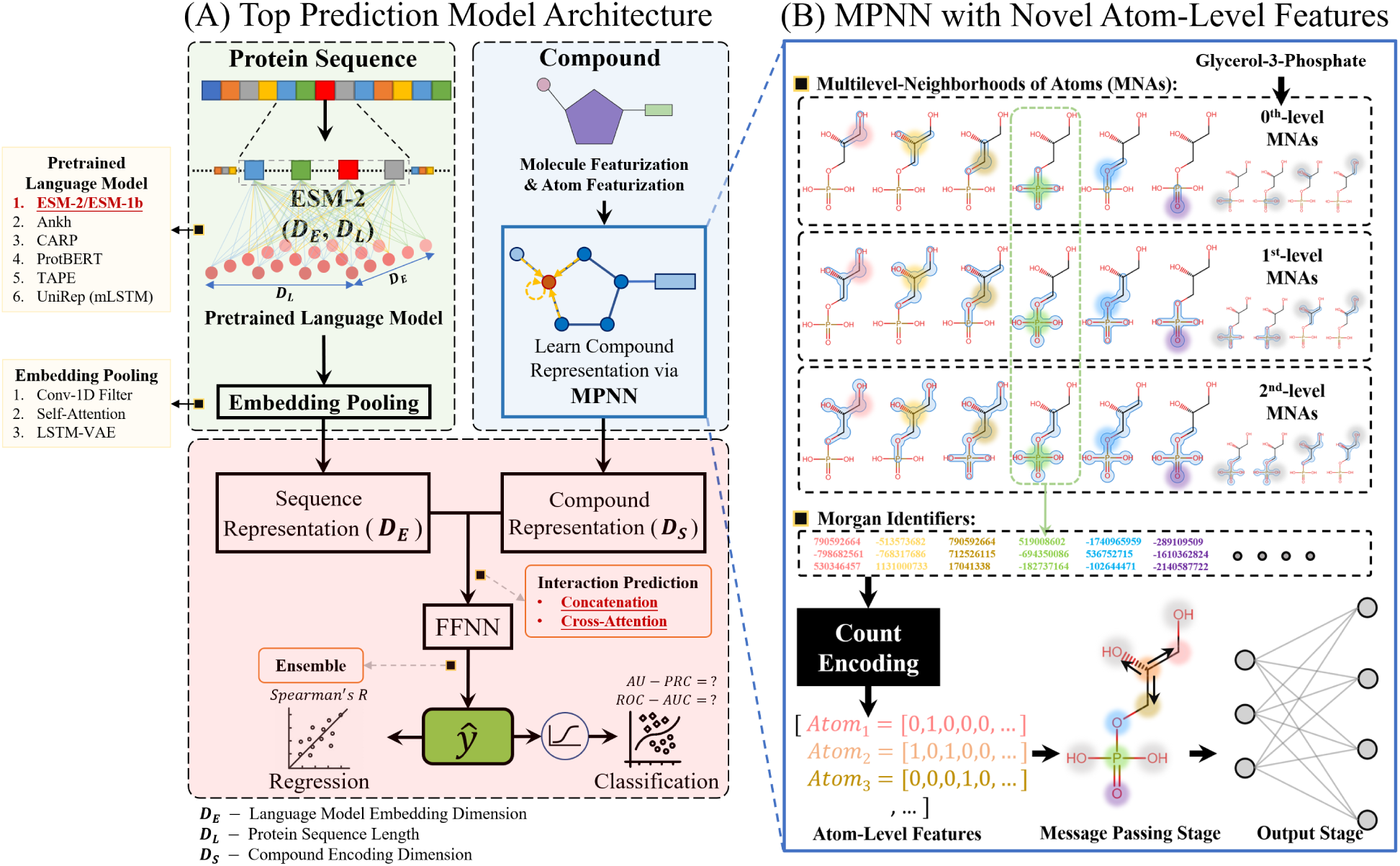
(A) Top model architecture for compound-protein interaction prediction. The main prediction pipeline contains three blocks, including, (1) sequence processing block (green) which generates and pools the sequence embeddings, (2) compound encoding block (blue) which generates compound representations and (3) interaction block (red) which learns the interactions between proteins and compounds. (B) Novel atom-level featurization of Glycerol-3-Phosphate (shows as an example). Multilevel neighborhoods of atoms (MNAs) substructures are identified surrounding each non-hydrogen atom before being converted to a set of Morgan fingerprint identifiers. Substructures surrounding each non-hydrogen atom are used as atom-level features and are then converted to numeric representations through count encoding (each dimension in the vector represents the count of occurrence of a different Morgan fingerprint identifier). The count-encoded substructures of molecule are input to the message passing stage of the MPNN, and further converted to the molecule encodings by the output stage.

### Processing of Kinetic Parameter Data

Approximately 60% of the kinetic parameters extracted from the BRENDA database ^20^ do not have annotated UniProt IDs for enzymes, which means there is no direct link between the functional data and the enzyme sequence. To address this issue, we leverage the recently developed KinMod database. KinMod^22^ encompasses the metabolic regulation network of 9814 organisms and employs a hierarchical data structure to signify relationships between proteins and kinetic information (obtained through *in vitro* experiments), with an emphasis on the small molecule regulatory network (SMRN). The hierarchical ontology of the KinMod database provides a novel framework that helps to retrieve, evaluate, and compare the functional metabolism of species and discover the extent of available and missing experimental values of metabolic regulation. Hence, it provides a standard framework for representing the available experimental data on enzyme kinetics and functions as a gap-filling tool for dealing with missing information in the metabolic network of various organisms. KinMod is therefore utilized to identify protein sequences based on EC number and species for those BRENDA entries with missing UniProt IDs. We create “core datasets” containing instances with labeled UniProt IDs in BRENDA, and create “pangenomic datasets” by augmenting the core datasets with KinMod-assigned sequences for data points with no UniProt annotation. To ensure data quality, pangenomic datasets are further screened using rigorous filters. Table 1 contains details of all the datasets retrieved and processed from BRENDA^20^. Supplementary Figure S1 illustrates the data processing steps involved in obtaining the core and pangenomic kinetic parameter datasets. A comparative t-SNE visualization and distributions of sequence lengths and target kinetic parameter values between the core and pangenomic datasets are provided in Supplementary Figure S2.

**Table 1:**
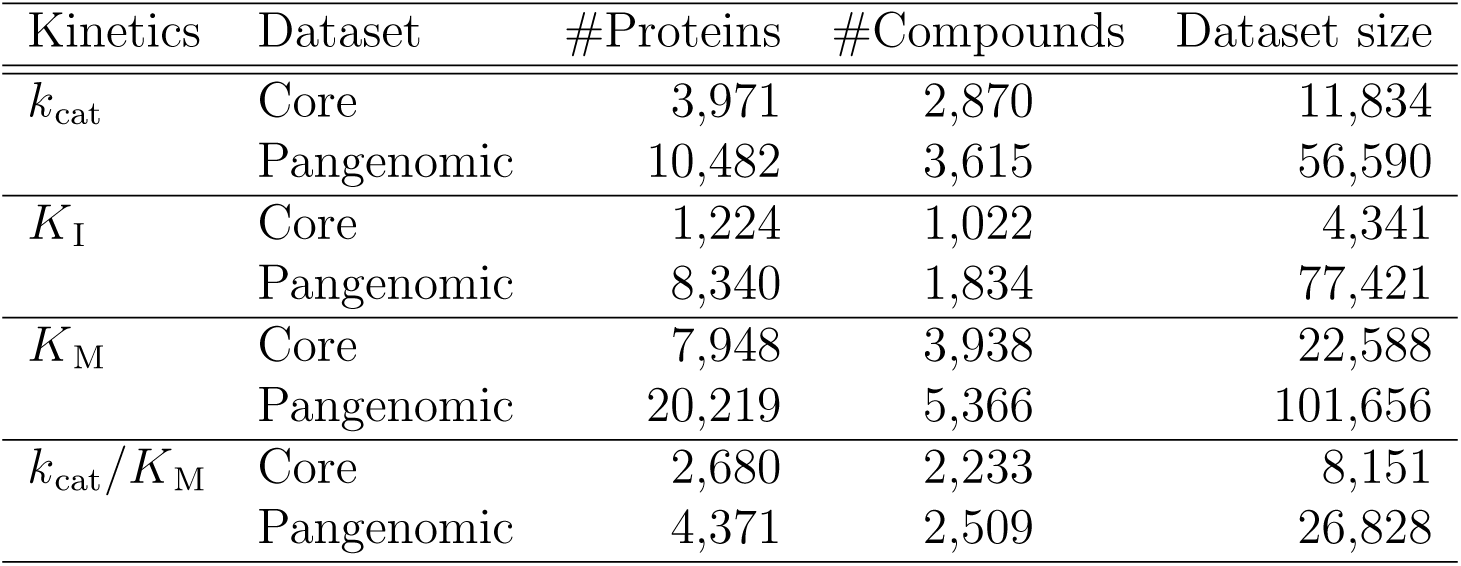
Size of each machine-learning-ready kinetic parameter dataset shown with number of sequences and compounds.

| Kinetics | Dataset | #Proteins | #Compounds | Dataset size |
| --- | --- | --- | --- | --- |
| $k_{\text{cat}}$ | Core | 3,971 | 2,870 | 11,834 |
|  | Pangenomic | 10,482 | 3,615 | 56,590 |
| $K_{\text{I}}$ | Core | 1,224 | 1,022 | 4,341 |
|  | Pangenomic | 8,340 | 1,834 | 77,421 |
| $K_{\text{M}}$ | Core | 7,948 | 3,938 | 22,588 |
|  | Pangenomic | 20,219 | 5,366 | 101,656 |
| $k_{\text{cat}}/K_{\text{M}}$ | Core | 2,680 | 2,233 | 8,151 |
|  | Pangenomic | 4,371 | 2,509 | 26,828 |

### Language Models and Sequence Embedding Pooling

The proposed prediction pipeline in this work starts with generating sequence embeddings for enzymes using protein language models (pLMs) pre-trained on massive protein sequence data. Four protein language models, including ESM-2^27^, ProtBERT^5^, Ankh^7^ and CARP^8^ are utilized in this study. We compare the performance of these pLMs on sequence-to-function learning tasks which we used to evaluate the robustness of different embedding pooling methods. Predictive performance of the compound-protein interaction prediction pipeline is mainly evaluated with ESM-2^27^ representations, while embeddings generated by other pLMs are also used to validate the proposed methods.

Sequence embeddings are very large (*d* × *L*, where *d* is the embedding dimension and *L* the sequence length) and thus require dimensionality reduction strategies that conserve as much information as possible from the embeddings. The traditional approach of average-pooling does not preserve the position-specific information of features in the original data, which is particularly important in sequence data like text or protein sequences. In our previous work^9^, we introduced a 1D convolutional pooling method and reported improved enzyme activity prediction. However, this method may not be optimal for datasets with large variations in sequence lengths. Convolutional pooling treats the entire sequence as a unit, and when sequences are padded to match the maximum sequence length in the data batch, the padded areas, carrying no meaningful information, can potentially dilute the information from shorter sequences. To address this limitation, we propose two alternative methods: self-attention pooling and LSTM variational encoder pooling.

Self-attention pooling is a novel strategy based on the scaled dot-product attention mechanism in the encoder block of the transformer. It employs a single attention head, transforming the unpadded sequence embeddings of varying shapes into a fixed-length vector with the same dimensionality as the original embeddings, maintaining the richness of the information while making it manageable for further processing.

The LSTM variational encoder pooling, on the other hand, uses an encoder layer to compress variable-length sequence embeddings into a consistent latent space vector. LSTM is capable of masking all the padding introduced to handle sequences of different lengths. The output from this architecture is normalized, enabling the LSTM encoder’s application beyond the training data distribution.

### Compound Encoding via MPNN with Novel Atom-level Featurization

Directed-message passing neural network^28^ (d-MPNN) is a recent compound property prediction framework that uses an improved message passing algorithm (i.e., based on messages associated with directed edges) and outperforms other existing models (i.e., MoleculeNet^25^) on 19 datasets of different compound properties. We incorporate this approach into our prediction pipeline to train a learnt molecule representation instead of the previously fixed representation based on chemical descriptors.

The d-MPNN has used basic atom properties (i.e., atom numbers, atom mass, etc.) as inputs to the neural network. Since we are training a relatively larger model (which contains a sequence processing block and interaction prediction block) than d-MPNN but with a relatively smaller dataset, we replace the default atom-level features with substructures surrounding each atom (as shown in Figure 1B). By introducing such prior structural information into the MPNN, we observe that the prediction performance is improved.

### Cross-attention for Learning Compound-Protein Interactions

Contemporary CPI prediction methodologies commonly employ a two-stage process, wherein protein and compound representations are generated and concatenated, followed by using a feed-forward neural network (FFNN) for quantitative target prediction. This approach may not adequately capture complex compound-protein interactions, as the feed-forward network’s role in learning these interactions is disproportionately small relative to the heavy architectures used for encoding sequences and molecules. Given the typically limited data available for CPI prediction, this can lead to a potential over-fitting risk. To address these limitations, we propose a cross-attention-based CPI prediction model architecture. The cross-attention mechanism allows the model to align and combine input representations, focusing on the most important parts, thereby effectively learning complex relationships^29^. This approach allocates a larger portion of learnable parameters to modeling the compound-protein interaction, thereby enhancing prediction performance.

### Ensembling Predictors

We adopt an ensemble approach to enhance prediction accuracy by aggregating outputs from four distinct models, each integrating different architecture combinations, namely: (1) MPNN-learnt compound representations with self-attention sequence pooling, (2) MPNN-learnt compound representations with LSTM sequence pooling, (3) MPNN-learnt compound representations with convolutional sequence pooling, and (4) cross-attention of compound substructure descriptors and sequence embeddings. The equal-weight ensemble prediction aims to capture a wider range of data features and patterns, leveraging the diverse strengths of each model.

### Data Splitting

In addition to examining the basic performance of the model by employing 5-fold cross-validation across the dataset, we further refine our evaluation through two focused tasks: the “protein design task” and the “compound design task”. For the protein design task, we utilize CD-HIT to cluster protein sequences, conducting experiments at both 80% and 60% similarity thresholds. These two levels of sequence identity introduce varying degrees of difficulty in sequence design tasks, aiming to test the model’s robustness in recognizing diverse protein functionalities. We implement 5-fold cross-validation by distributing the sequence clusters obtained from CD-HIT equally among the five folds, ensuring comprehensive evaluation across the entire dataset.

In the compound design task, compounds are clustered based on structural fingerprints and Tanimoto similarity. Clustering is performed using similarity thresholds of 0.2 and 0.4, representing two levels of difficulty in the design of chemical compounds, ensuring a rigorous test of the model’s predictive capabilities. Each compound is uniquely assigned to a cluster, thereby preventing any overlap across the five folds in our 5-fold cross-validation setup. This methodological choice guarantees that each fold presents a structurally diverse set of compounds, enhancing the model’s ability to predict kinetic parameters for compounds not similar to those previously encountered during training.

By using 5-fold cross-validation in both the protein and compound design tasks, we obtain a more reliable prediction performance evaluation across the entire dataset, thereby advancing our understanding in protein/enzyme discovery and enzyme engineering.

### Model Architecture Blocks Tests

The performance of different protein sequence encoding methods as well as compound encoders was studied through the design of a set of tests for the CPI-Pred model. The improved compound encoding MPNN, incorporating novel atom-level featurization, was tested against other encoding methods like d-MPNN, ECFP6 count encoding, and Morgan fingerprint bit vector representations on *k*_cat_ and *K*_I_ core datasets, as further discussed in the Supplementary Information. Concurrently, in a study testing the efficacy of different sequence encoding blocks, we assessed various encodings across *k*_cat_ and *K*_M_ datasets, including *n*-gram encoding, sequence embeddings with traditional pooling strategies, and the novel self-attention pooling method.

### Design of the Error Predictor

To enhance the reliability of our compound-protein interaction model, we integrated an error predictor designed to assess the confidence level of the predictions. This error predictor’s architecture calculates Euclidean distances for both protein sequences and compounds from a novel query against all entries in the model’s training dataset. Distances for protein sequences are derived using embeddings generated by the protein language model ESM-2, while distances for compounds are computed using Extended Connectivity Fingerprints (ECFP6) count encodings. These encodings are used because they directly encode molecule substructure without needing to learn those representations (MPNN), therefore reflecting the distances between the structural information of the compounds better. Both protein sequence distances and compound distances are associated with absolute error values derived from the CPI-Pred model (the deviation between predicted interaction values and actual observed values for each data point in the training set). The absolute errors are classified as either ones of high error if the error indicates a difference of about 2 orders of magnitude between the predicted and actual values, and low error if the error indicates an error of less than 2 order of magnitude between the predicted and actual values. We organize these distance-error pairs into 100 percentile groups for both proteins and compounds, thereby forming a detailed distribution profile for each category. Within each percentile group, we compute the mean, standard deviation, maximum, and minimum of the distances and errors. These statistics well characterized the distribution of distances between the novel query and the entire training set of the predictive model. The aggregated statistics are then used as input features to train a three-layer neural network. This training equips the neural network with the capability to classify the reliability of new predictions into high or low error intervals based on the learned error distributions. This systematic approach ensures that the error predictor enhances the reliability and interpretability of the compound-protein interaction predictions.

## Results and Discussion

In this section, we systematically evaluate our CPI prediction model and its key components for modeling proteins and compounds: sequence encodings (including pLM embeddings and pooling methods) and compound encodings. We compare different protein language models by assessing sequence-to-function learning on two experimental datasets: the PafA^17^ point mutations dataset and the avGFP dataset from Sarkisyan^23^. In addition to compound encoder comparisons, we perform a study to test the impact of various sequence encoding techniques (including one-hot encoding, sequence embeddings with and without pooling strategies and attention mechanisms) on the model’s performance across *k*_cat_ and *K*_I_ datasets. The performance of different compound encoding methods, including the improved MPNN-learnt representation used in CPI-Pred, original d-MPNN compound encodings, and chemical descriptors, is assessed on *k*_cat_ and *K*_I_ core datasets. Lastly, we benchmark the predictive performance of CPI-Pred against baseline and state-of-the-art models on ten datasets (summarized in Supplementary Table S1) which cover a diverse range of compound-protein interactions such as kinetic property prediction, enzyme activity prediction, and binding affinity prediction. This represents, to our knowledge, the most comprehensive testing in the domain of protein engineering and kinetic property prediction and is available for the community to benchmark any future methods.

### Sequence Embeddings and Pooling Methods

We compare the effectiveness of four protein language models (ProtBERT^5^, ESM-2^27^, CARP^8^ and Ankh^7^) in our pipeline by evaluating sequence-to-function learning on the two experimental datasets (PafA and avGFP) mentioned above. The PafA dataset consists of 1034 point mutations of PafA sequence (modifying or deleting one single amino acid in each sequence variant) and corresponding *k*_cat_ values, and the avGFP^23^ (Aequorea victoria green fluorescent protein) dataset comprises 54,000 sequences from the local fitness landscape of avGFP.

Based on the performances of sequence-to-function learning with results detailed in Supplementary Table S3, we determined that ESM-2 consistently outperformed the other embedding models, thereby emerging as the best choice for our CPI prediction model. Moreover, the self-attention-based along with the 1-D convolutional pooling methods excelled over other embedding pooling methods, achieving the highest performance in sequence-to-function learning (Supplementary Table S2 & Supplementary Table S3).

This conclusion is further validated by the results highlighted in Supplementary Table S6, which shows that a model with the self-attention pooling method is able to match the performance of a model that uses no pooling method for the sequence embeddings, while allowing for a model that is much smaller in size. The superior predictive performance of the ESM-2 language model and self-attention pooling in these two tasks supports their promising applicability in CPI prediction tasks.

### Compound Encoders

An overall good predictive capability of the improved MPNN-learnt compound representation (used by CPI-Pred) is shown in Supplementary Table S5, where predictions are tested on *k*_cat_ and *K*_I_ core datasets over different molecule encoding methods, including (1) improved MPNN-learnt representation, (2) directed-message passing neural network (d-MPNN) with default settings found in the original paper, (3) ECFP6^30^ count encoding method used in Xu et al.^9^, and (4) Morgan fingerprint^31^ bit vector representation commonly used in existing CPI prediction models. The MPNN-learnt compound representation consistently outperforms other methods with statistically significant improvements.

### CPI-Pred’s Performance Against State-of-the-art CPI Models

Here, we showcase the performance of our CPI-Pred model through an extensive comparative analysis. In particular, we benchmark our model’s performance against DLkcat, a previously developed state-of-the-art method for kinetic property prediction and a robust KNN baseline. The prediction tasks we consider encompass a broad spectrum:

1. The four aforementioned kinetic parameters (*k*_cat_, *K*_I_, *K*_M_, and *k*_cat_*/K*_M_) from BRENDA.
2. Experimental enzyme activities data, including phosphatase activities ^32^, kinase-ligand binding affinities ^33^, halogenase activities ^34^, esterase hydrolytic activities ^35^, and amino-transferase activities ^36^.
3. PDBbind binding affinity dataset, which has been extensively studied.

Figure 3, Figure 2 and Figure 4 illustrates correlations between predicted values and experimental measurements across this diverse array of prediction tasks.

**Figure 2:**
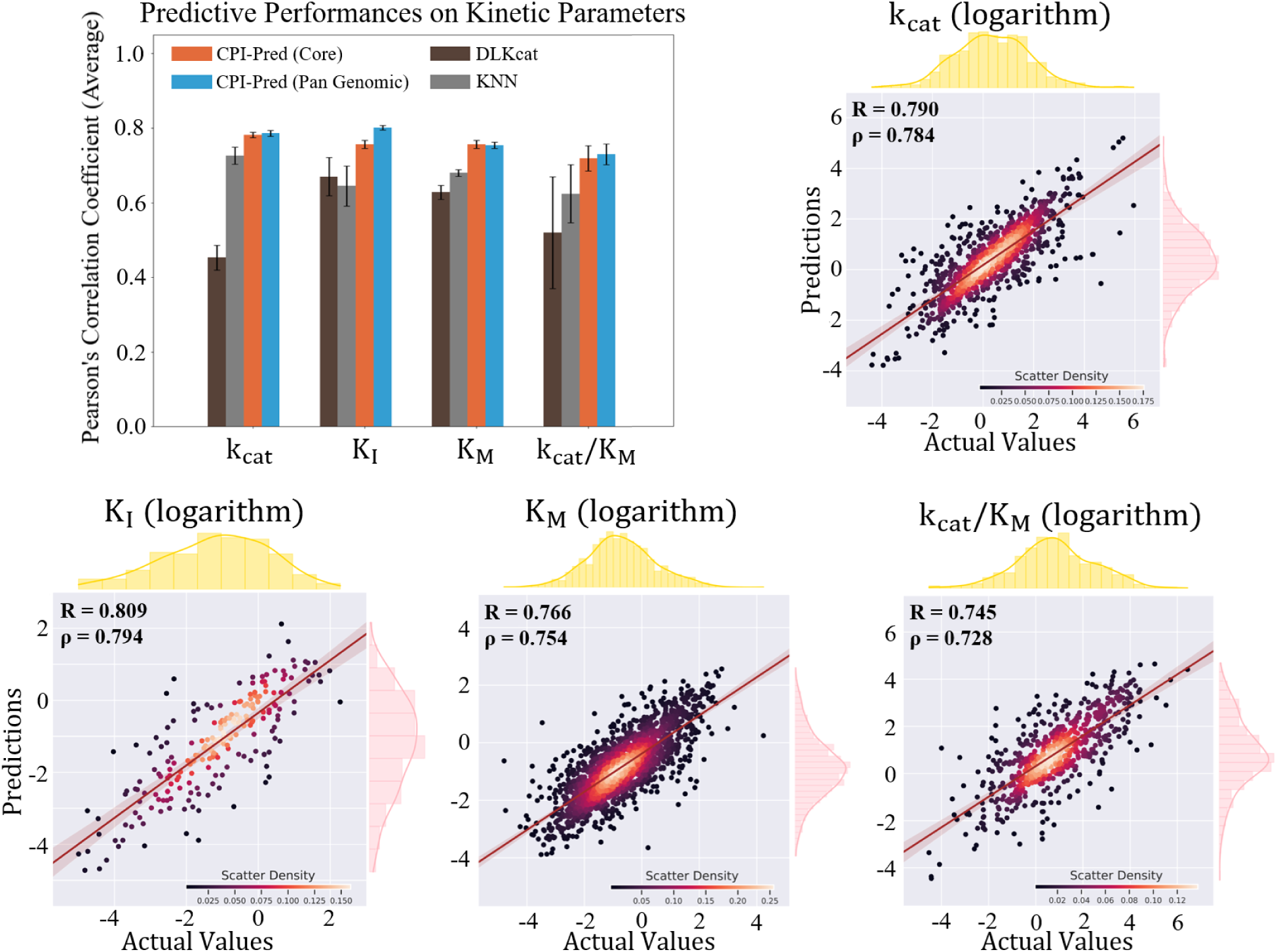
Comparison of CPI-Pred’s performance with other models on kinetic parameters datasets. The barplot shows the performance of CPI-Pred, DLkcat, and KNN models across four datasets: *k*_cat_, *K*_I_, *K*_M_, and *k*_cat_*/K*_M_ (all values logarithmic). Models are colored similarly to the previous figure: brown for DLkcat, grey for KNN, orange for CPI-Pred trained on the core dataset, and blue for CPI-Pred trained on the pangenomic dataset. Below the barplot, scatter plots illustrate predicted versus actual values for each kinetic parameter. Pearson’s *r* and Spearman’s *ρ* are displayed for each scatter plot, with adjacent histograms for actual values (yellow) and predicted values (pink).

**Figure 3:**
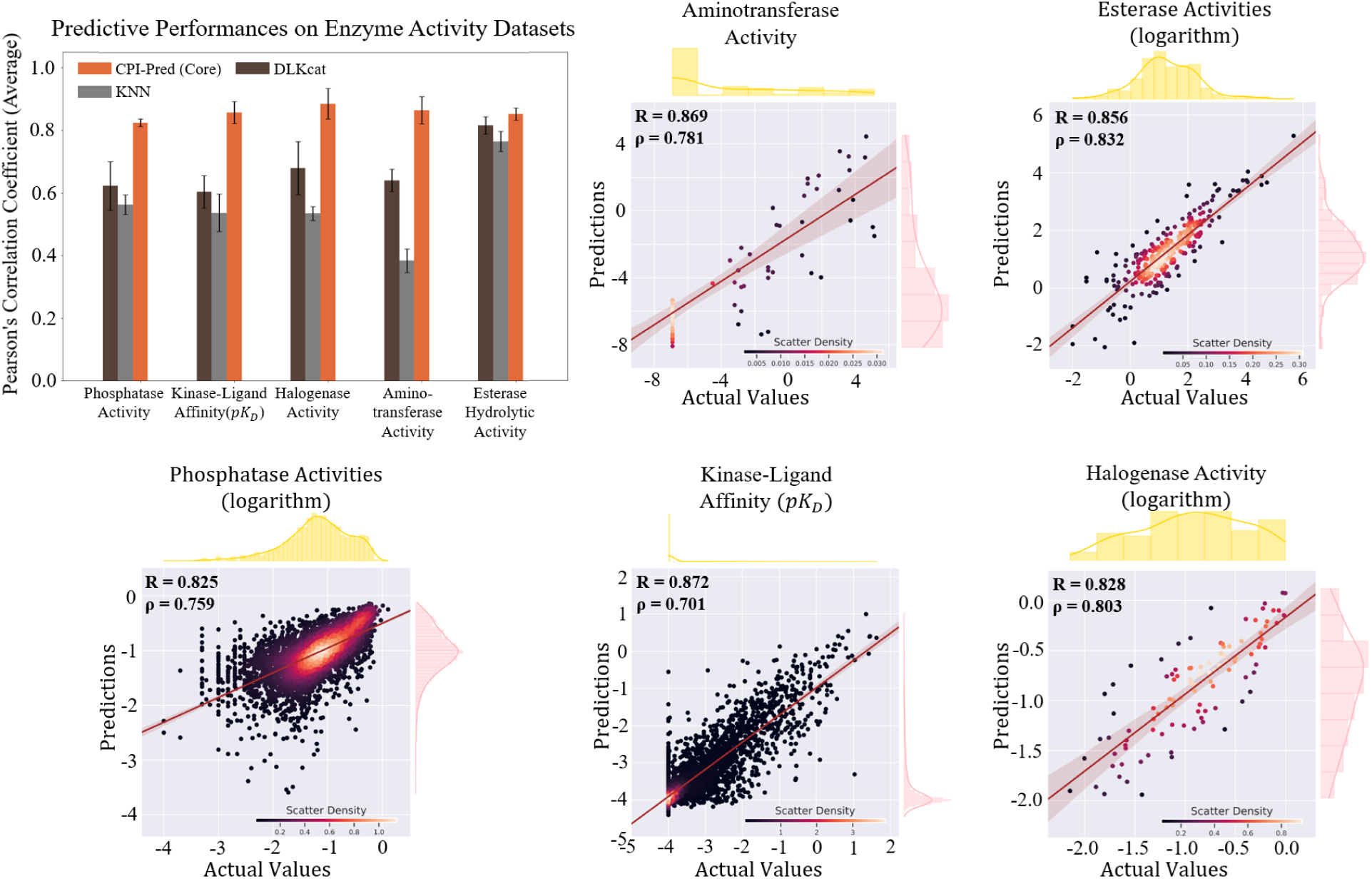
Comparison of CPI-Pred’s performance with other models on five experimental enzyme activity datasets. The barplot shows the performance of CPI-Pred, DLkcat, and KNN models across the datasets. Models are denoted by distinct colors: brown for DLkcat, grey for KNN, and orange for CPI-Pred trained on the core dataset. The metric used is Pearson’s correlation coefficient, and error bars indicate the standard deviation across 5-fold cross-validation. Below the barplot, scatter plots show predicted versus actual values for each enzyme activity dataset: *k*_cat_, aminotransferase activities, esterase hydrolytic activities, phosphatase activities, kinase ligand affinities, and halogenase activities. Pearson’s *r* and Spearman’s *ρ* are provided in each plot, with adjacent distributions of actual values (yellow histograms) and predicted values (pink distributions).

**Figure 4:**
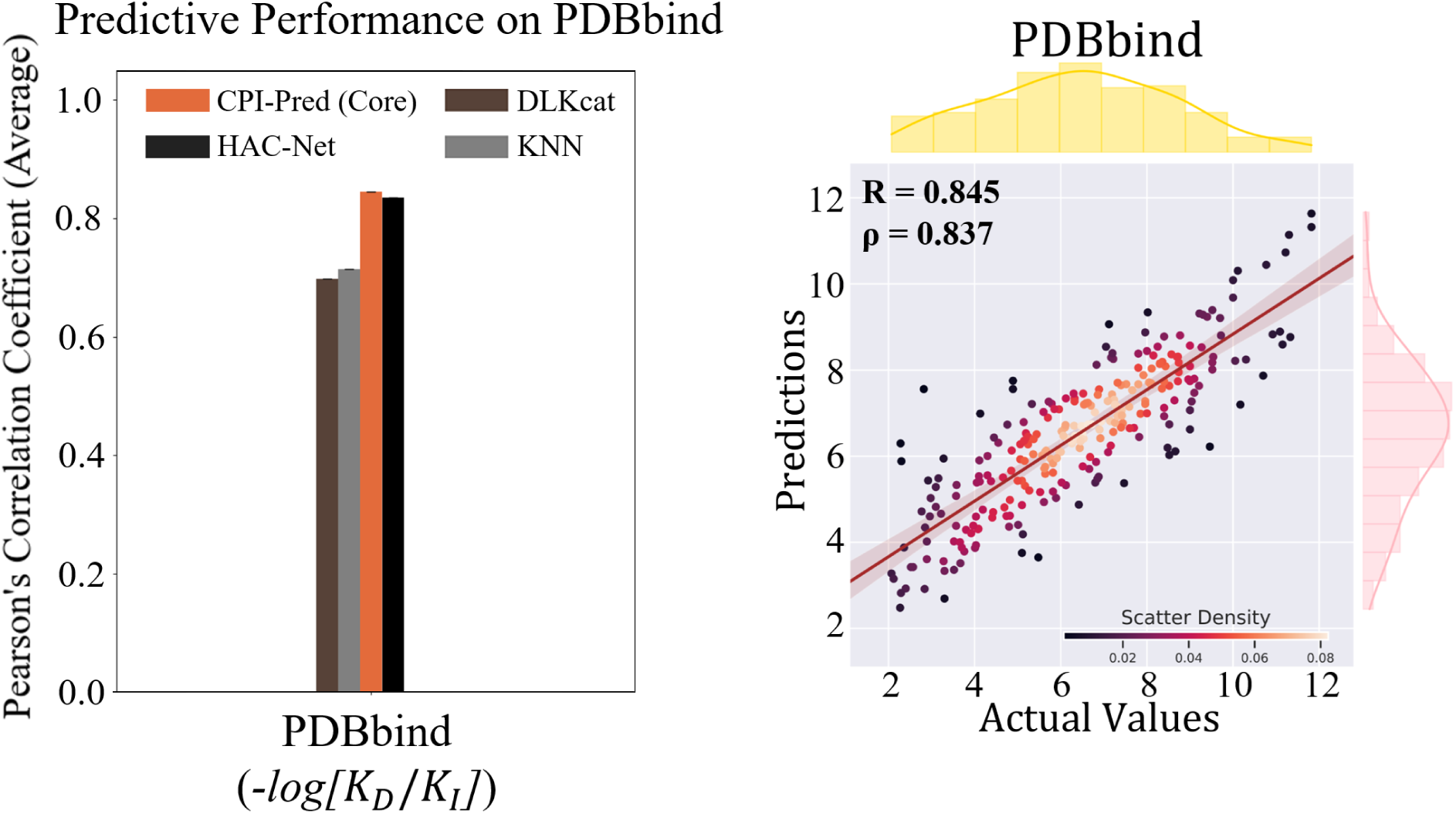
Comparison of CPI-Pred’s performance with other models on the PDBbind dataset. The barplot shows the performance of CPI-Pred, DLkcat, and KNN models, along with the HAC-Net model (denoted by the black bar). The metric used is Pearson’s correlation coefficient, with error bars indicating standard deviation across 5-fold cross-validation. Below the barplot, a scatter plot shows predicted versus actual values for the PDBbind dataset. Pearson’s *r* and Spearman’s *ρ* are displayed in the scatter plot, with adjacent distributions of actual values (yellow histogram) and predicted values (pink distribution).

#### Predicting kinetic parameters

For each of the four kinetic parameters, we present the performance of the CPI-Pred model trained on the core dataset and on the pangenomic dataset (Figure 2).

All test sets for bench-marking kinetic parameter predictions are drawn from their corresponding BRENDA core datasets. Across the board, the CPI-Pred model trained on the pangenomic dataset surpasses the KNN, DLkcat, and the CPI-Pred model trained on the core dataset (Figure 2). Additionally, CPI-Pred achieves a higher performance relative to recent state-of-the-art models such as *K*_M_ Prediction by Kroll et al.^13^ and TurNuP^37^.

These models also reported predictions on kinetic parameter datasets, sourced from BRENDA and processed in a manner similar to our core dataset. Moreover, CPI-Pred excels in both the “protein design” and “compound design” tasks (as shown in Table 2), which demonstrate its ability to predict properties related to unseen compounds and unfamiliar sequences.

**Table 2:** Prediction performance (Pearson’s correlation coefficients) of CPI-Pred on compound and protein design tasks of kinetic parameters, with KNN and DLkcat used for bench-marking. Only core datasets were used for testing. The shown results correspond to averages and standard deviations across models trained using 5-fold cross-validation. The Tanimoto coefficient^38^ is used to set the similarity threshold, meaning a Tanimoto coefficient of 1 means that 2 compounds have identical structural descriptors. CD-HIT^39^ is used to split the sequences basd on sequence similarity.

| Tasks | Method | $k_{\text{cat}}$ | $K_{\text{I}}$ | $K_{\text{M}}$ | $k_{\text{cat}}/K_{\text{M}}$ |
| --- | --- | --- | --- | --- | --- |
| Simple Task<br>Random Split | CPI-Pred (Pangenomic) | 0.786±0.008 | 0.801±0.006 | 0.754±0.009 | 0.730±0.028 |
|  | CPI-Pred (Core) | 0.781±0.005 | 0.756±0.011 | 0.756±0.011 | 0.719±0.034 |
|  | KNN | 0.726±0.023 | 0.645±0.054 | 0.680±0.009 | 0.624±0.078 |
|  | DLkcat | 0.453±0.033 | 0.670±0.051 | 0.628±0.019 | 0.520±0.051 |
| Compound Design<br>(Similarity<br>Threshold: 0.2) | CPI-Pred (Pangenomic) | 0.747±0.008 | 0.728±0.032 | 0.720±0.028 | 0.620±0.049 |
|  | CPI-Pred (Core) | 0.725±0.006 | 0.630±0.035 | 0.697±0.036 | 0.612±0.045 |
|  | KNN | 0.670±0.028 | 0.536±0.056 | 0.581±0.038 | 0.545±0.046 |
|  | DLkcat | 0.390±0.039 | 0.491±0.056 | 0.546±0.050 | 0.425±0.052 |
| Compound Design<br>(Similarity<br>Threshold: 0.4) | CPI-Pred (Pangenomic) | 0.730±0.049 | 0.712±0.019 | 0.653±0.037 | 0.601±0.030 |
|  | CPI-Pred (Core) | 0.615±0.054 | 0.596±0.047 | 0.607±0.043 | 0.582±0.027 |
|  | KNN | 0.571±0.034 | 0.487±0.071 | 0.471±0.060 | 0.480±0.055 |
|  | DLkcat | 0.435±0.052 | 0.510±0.064 | 0.464±0.035 | 0.404±0.052 |
| Protein Design<br>(Sequence<br>Identity: 80%) | CPI-Pred (Pangenomic) | 0.527±0.071 | 0.673±0.052 | 0.653±0.040 | 0.396±0.047 |
|  | CPI-Pred (Core) | 0.536±0.050 | 0.662±0.043 | 0.648±0.051 | 0.392±0.050 |
|  | KNN | 0.380±0.071 | 0.493±0.011 | 0.516±0.036 | 0.325±0.042 |
|  | DLkcat | 0.407±0.056 | 0.644±0.064 | 0.577±0.052 | 0.366±0.038 |
| Protein Design<br>(Sequence<br>Identity: 60%) | CPI-Pred (Pangenomic) | 0.472±0.050 | 0.648±0.050 | 0.602±0.052 | 0.387±0.059 |
|  | CPI-Pred (Core) | 0.498±0.046 | 0.610±0.058 | 0.528±0.048 | 0.399±0.050 |
|  | KNN | 0.396±0.040 | 0.405±0.038 | 0.476±0.056 | 0.377±0.042 |
|  | DLkcat | 0.407±0.056 | 0.532±0.053 | 0.497±0.052 | 0.401±0.042 |

#### Predicting enzyme activities

In the case of enzyme activities datasets, our CPI-Pred model demonstrated superior performance, achieving consistently and substantially better results than DLkcat and KNN models (Figure 3). This result highlights the robustness of our model in accurately predicting enzyme activities.

#### Predicting binding affinity

The last dataset evaluated was PDBbind, with the CPI-Pred model trained on the PDBbind 2020 version and tested on the frequently cited PDBbind 2016 core dataset^25^, which consists of approximately 200 structures. The PDBbind database is a comprehensive collection of experimentally measured binding affinities for biomolecular complexes, providing 3D Cartesian coordinates of both ligands and their target proteins, thereby allowing for structure-based featurization. CPI-Pred’s performance was compared with HAC-Net, which to the best of our knowledge, has hitherto achieved the best predictions on the PDBbind 2016 core dataset. Strikingly, CPI-Pred achieved a correlation of 0.845, slightly surpassing HAC-Net’s correlation of 0.835 (Figure 4); both methods performed substantially better than DLkcat and KNN models. This result is all the more remarkable given that CPI-Pred, unlike HAC-Net and most other binding affinity prediction models, does not require binding pocket information as input. This aspect of CPI-Pred broadens the model’s applicability, particularly for unknown or novel sequences in protein engineering.

Taken together, the above results demonstrate that CPI-Pred can deliver top-tier performance while utilizing a more constrained set of input information, underscoring its potential for broader utility in the field.

### Error Predictor Performance

While we have shown CPI-Pred is valuable for property prediction, the accuracy is dependent on the quality of the dataset, especially for kinetic property predictions. Hence, as described earlier, we sought to develop an additional element of the pipeline to also predict the error level of the CPI-Pred predictions using an error predictor pipeline. The efficacy of the error predictor was evaluated through its application to the *k*_cat_ dataset. This dataset consists of a diverse array of compound-protein interactions. The error predictor achieved a classification accuracy of 71%, demonstrating a substantial capability of accurately discern between high and low confidence. High and low confidence classes were defined based on the absolute error between the log-transformed predicted and true values, with a threshold of 0.2 used to categorize the predictions.

This accuracy highlights the predictor’s effectiveness in leveraging training set error patterns to forecast the reliability of predictions on new data. The error predictor enhances confidence in the interaction model’s outputs by significantly reducing the risk of making potentially inaccurate predictions. Additionally, it can be helpful when designing subsequent experimental validations and collecting more data points by focusing on those landscapes identified with low confidence, thereby contributing to the more comprehensive completion of the compound-protein interaction landscape. For example, the error prediction in Supplementary Figure S5 highlights the relative lack of kinetics data in enzyme class 4 (lyases), suggesting the need for additional characterization of enzymes in this class.

## Limitations

While CPI-Pred performs well on relatively smaller enzyme activity datasets, its predictive accuracy declines when tasked with predicting novel sequences and compounds. Applying a pangenomic approach, as we did for kinetic parameters from BRENDA, is challenging for these enzyme activity datasets, particularly when novel sequences or compounds are involved. This makes the need for high-quality, high-fidelity data even more crucial, yet the acquisition of such data via experimentation is costly. Furthermore, the BRENDA kinetics data that we utilized carries uncertainties due to variable experimental conditions such as pH, temperature, buffers used, and other potential human errors, which may affect predicted kinetic parameters. Despite our efforts to mitigate this by filtering out instances significantly affected by experimental conditions and through the use of an error predictor module, this uncertainty remains a limitation inherent to the available data.

## Conclusion

In conclusion, the development of deep learning techniques can revolutionize the systematic engineering of proteins, microbial strains and communities and advance the fields of systems and synthetic biology. In particular, the area of predicting protein-related properties and functions from sequence can be accelerated by leveraging advances in the development of large language models. In this paper, we have presented a novel deep learning-based workflow for CPI prediction, which utilized updated protein language models, improved sequence embedding pooling algorithms, learnt representation of compounds via a message-passing neural network with customized atom-level features, and a cross-attention block for better interaction prediction. We have also demonstrated the utility of this model architecture by showing that it achieves good performance with different embeddings and outperforms other existing state of the art models and non-neural network baselines on a variety of CPI datasets. Finally, the CPI datasets we have compiled are comprehensive and will be valuable to benchmark future CPI prediction algorithms and represent a significant milestone for this community. Our work provides a promising direction for future research in developing accurate and robust CPI prediction models and encourages the collection and utilization of high-quality data to drive the advancement of this exciting field.

## Supplementary Information

Additional model development details, including data scraping and processing and validation, model architecture details, model bench-marking tests set-up and results are all included in the attached SI.

## Acknowledgement

The authors thank Genome Canada, the Ontario Ministry of Research and Innovation, the Miller Institute, as well as the Mitacs program for their generous funding. The research of YSS is supported in part by an NIH grant R35-GM134922.

## Notes

https://github.com/Zhiqing-Xu/CPI-Pred

### Competing Interest Statement

The authors have declared no competing interest.

