## Supplemental Information for "CPI-Pred: A deep learning framework for predicting functional parameters of compound-protein interactions"

#### Processing of Kinetic Parameter Data

The data processing steps involved in obtaining the core and pangenomic kinetic parameter datasets are visually represented in Figure S1.

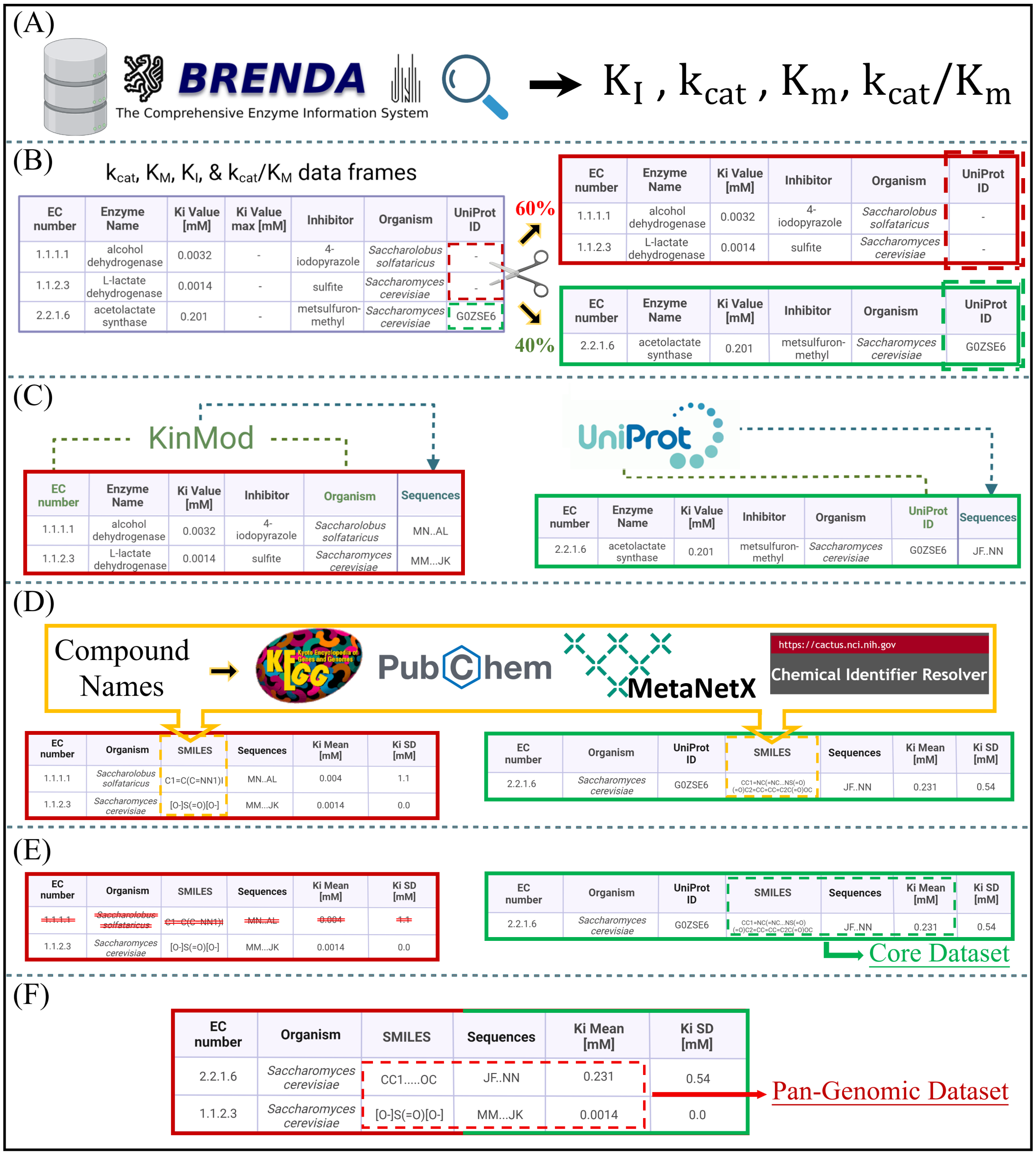


Figure S1: The figure presents six steps (A-F) in the processing of Kinetic Parameter Data. (For full caption see next page.)

Figure S1: (Continued) (A) The process begins with the extraction of kinetic datasets from BRENDA, which includes parameters such as enzyme turnover numbers, Michaelis-Menten constants, inhibition constants, and catalytic efficiency. (B) The data are then divided based on the availability of UniProt IDs: a dataset composed of instances with UniProt ID annotations, and another one containing instances without such information. (C) In the next step, sequences are incorporated into the datasets. For the dataset lacking UniProt IDs, sequences are obtained from KinMod using EC number and organism. (D) Compound names (not depicted in the figure) are cross-referenced in three databases (i.e., KEGG, MetaNetX, PubChem). Those compounds not found in these databases are searched using the Chemical Identifier Resolver (CIR) from NCI/NIH to obtain the SMILES representations. (E) The instances in each dataset are grouped by compound-sequence pairs, and the mean and standard deviation of kinetic values are computed. Any compound-sequence pair with a standard deviation exceeding a preset threshold is filtered out. (F) In the final step, the two datasets prepared in the previous steps are merged, and grouped by organism, EC number, and compound. The mean and standard deviation of kinetic values are again computed, with any organism-EC-compound group whose standard deviation exceeds the preset threshold being filtered out.

### Size of all datasets used to benchmark CPI-Pred

Table S1: Size of each machine-learning-ready kinetic parameter dataset shown with number of sequences and compounds.

| Interactions/Kinetics | Dataset | #Proteins | #Compounds | #Data | Reference |
|---|---|---|---|---|---|
| $k_{cat}$(Turnover Number) | Core | 3,971 | 2,870 | 11,834 | Chang et al.[1] |
| | Pangenomic | 10,482 | 3,615 | 56,590 | |
| $K_I$(Inhibition Constant) | Core | 1,224 | 1,022 | 4,341 | Chang et al.[1] |
| | Pangenomic | 8,340 | 1,834 | 77,421 | |
| $K_M$(Michaelis Constant) | Core | 7,948 | 3,938 | 22,588 | Chang et al.[1] |
| | Pangenomic | 20,219 | 5,366 | 101,656 | |
| $k_{cat}/K_M$(Catalytic Efficiency) | Core | 2,680 | 2,233 | 8,151 | Chang et al.[1] |
| | Pangenomic | 4,371 | 2,509 | 26,828 | |
| Catalytic Activity | Phosphatase | 218 | 168 | 36,624 | Huang et al.[2] |
| Catalytic Activity | Halogenase | 42 | 62 | 2,604 | Fisher et al.[3] |
| Catalytic Activity | Aminotransferase | 25 | 18 | 450 | Li et al.[4] |
| Catalytic (Hydrolytic) Activity | Esterase | 147 | 96 | 14,112 | Martínez-Martínez et al.[5] |
| Ligand Affinity | Kinase | 405 | 72 | 29,160 | Davis et al.[6] |
| Binding Affinity | PDBbind 2020 | 8,053 | 10,893 | 12,377 | Wang et al.[7] |
| | PDBbind 2016 Core | 192 | 263 | 263 | Wu et al.[8] |

### Validation of Pangenomic dataset

In order to validate the use of this pangenomic data processing approach, a t-SNE[9] plot was used as a visualization method to observe the compound-protein clusters and corresponding kinetic values of the core and pangenomic datasets. Furthermore, the distributions of the pangenomic and core datasets were compared to ensure that no hallucinated distributions were introduced by the data processing method. Figure S2(A) shows a tSNE plot that visualizes the data points from $K_I$ core dataset (shown by red/orange points) against $K_I$ pangenomic dataset (data points shown by blue purple points). This t-SNE plot serves as a justification of the pangenomic approach, which is demonstrated by the fact that most data points from core dataset are surrounded by (or at least close to) a data cluster from pangenomic dataset. This shows the similarities in the distributions of core vs pangenomic dataset. Figure S2(B) and Figure S2(C) present the violin plots comparing the core and pangenomic datasets for each of the three kinetic parameters. The t-SNE plot depicts consistent compound-protein clusters across the pangenomic and core datasets with respect to the output kinetic values, which suggests that the pangenomic approach does not hallucinate new and dissimilar input data clusters. Both the target value and sequence length distributions remain consistent for both the pangenomic and core datasets as well, with slightly higher peaks for the sequence lengths in the pangenomic dataset.

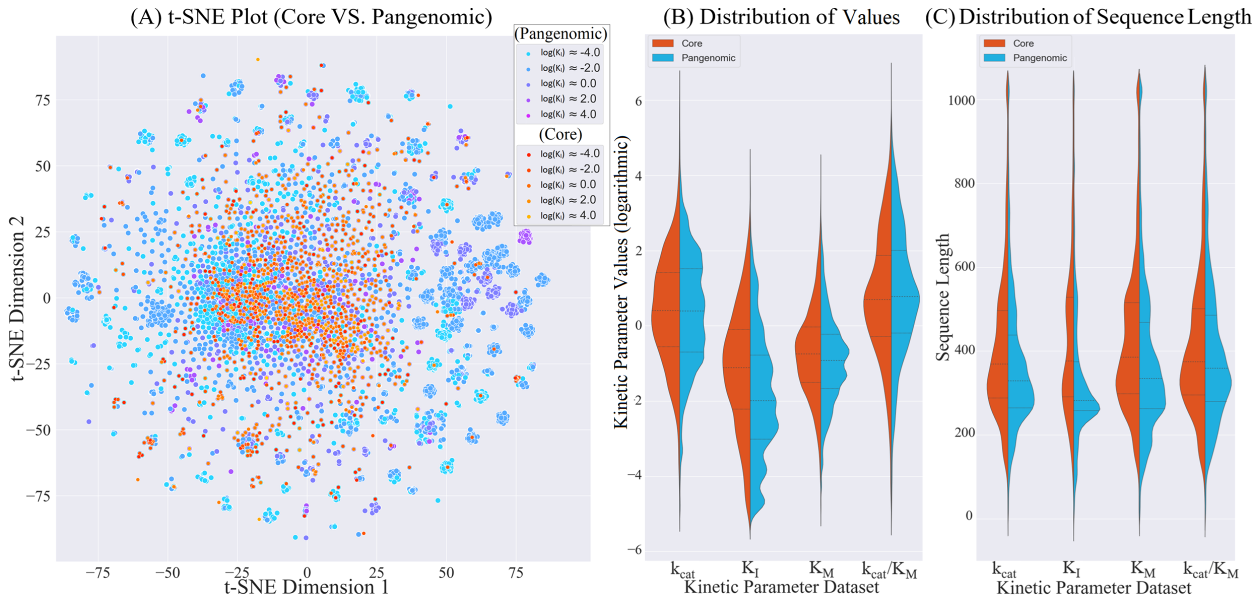


Figure S2: t-SNE and distribution plots comparing the core and pangenomic datasets. (A) A t-SNE plot that visualizes the data points from $K_I$ core dataset (shown by red/orange markers) and $K_I$ pangenomic dataset(shown by blue/purple markers), illustrating consistent compound-protein clusters for core and pangenomic datasets with respect to output kinetic values. (B) Distributions of target kinetic parameter values for (from left to right) $k_{cat}$ core, $k_{cat}$ pangenomic, $K_I$ core, $K_I$ pangenomic, $K_M$ core, and $K_M$ pangenomic datasets. The core datasets are shown by orange distributions while the pangenomic dataset are shown by blue distributions. (C) Sequence length distributions in the corresponding core and pangenomic datasets for $K_I$, $k_{cat}$, and $K_M$.

### Model Compound-Protein Interactions Using a Cross-attention Block

A cross-attention block is used to replace the commonly used simple concatenation of compound and sequence representations. Figure S3 shows how prediction is computed assuming one single attention head being used.

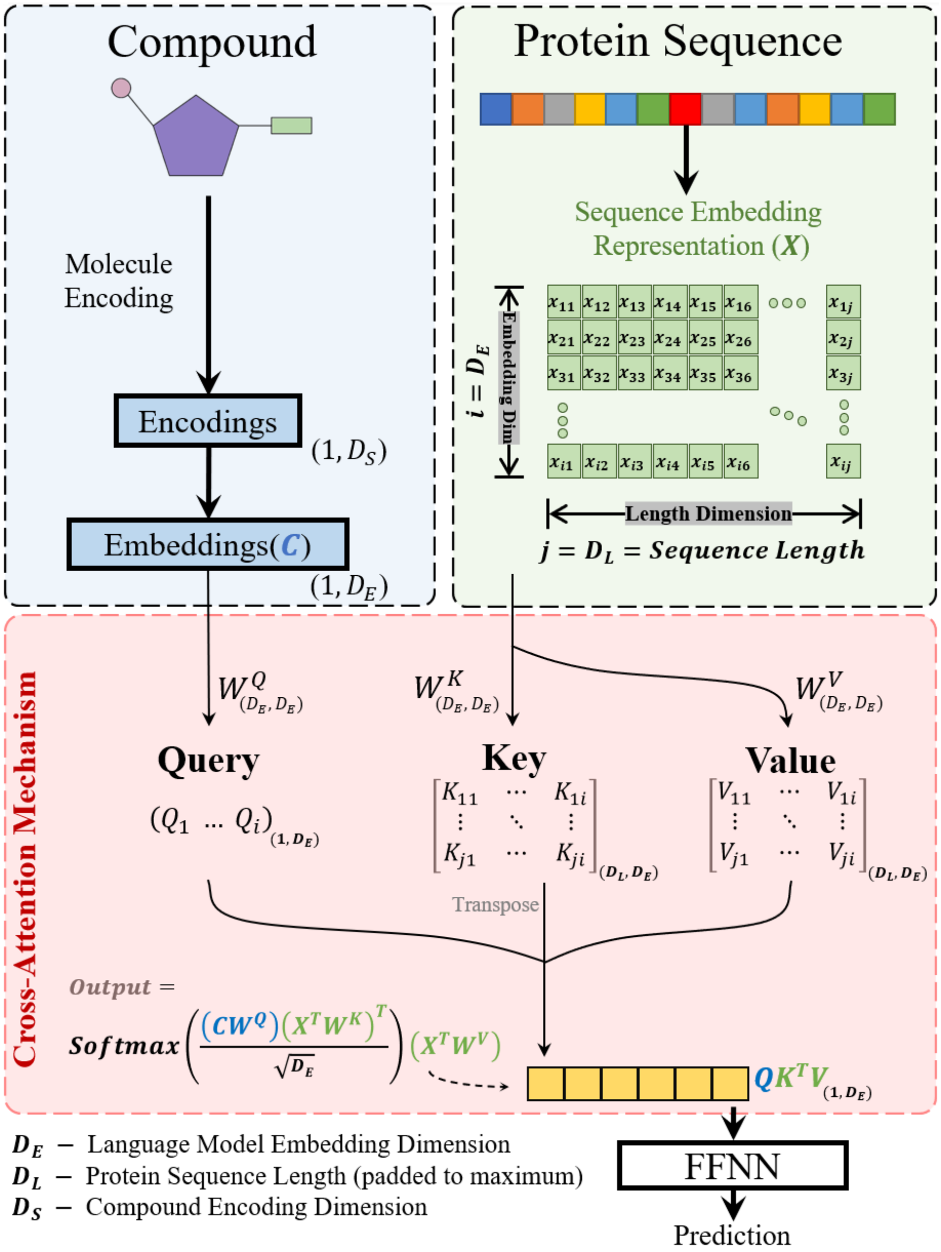


Figure S3: Schematic of the proposed cross-attention mechanism for compound-protein interaction prediction. A compound encoding of dimension (1, $D_S$) is first converted to the same dimensionality as the sequence embeddings ($D_E$) via a PyTorch embedding layer, yielding a compound embedding $C$ of dimension (1, $D_E$). The sequence embedding $X$, representing a protein sequence, has dimensions ($D_E$, $D_L$), where $D_L$ is the sequence length. These embeddings are transformed into queries (from the compound), and keys and values (from the sequence) via linear transformations: $C$ is multiplied by a weight matrix $W_Q$ to generate the query matrix (of dimension (1, $D_E$)), and $X$ is multiplied by weight matrices $W_K$ and $W_V$ to generate the key and value matrices (both of dimension ($D_L$, $D_E$)). The output of the cross-attention block is then computed as softmax$((CW_Q)(X^TW_K)^T/\sqrt{D_E})(X^TW_V)$, resulting in a matrix of dimension (1, $D_E$). This output is subsequently fed into a three-layer feed-forward neural network for final prediction, providing a prediction for the interaction between the given compound and protein sequence.

### Error Level Predictor Architecture

Figure ?? is a figure showing the architecture of the error level prediction for compound-protein interactions. It illustrates the process of calculating Euclidean distances for a new protein-compound pair relative to the training sets used in the CPI-Pred model. The distances are categorized into percentile ranks for proteins and compounds separately, with statistical measures (mean, standard deviation, maximum, and minimum) computed for each rank. These statistics form the inputs for a neural network, which is then trained to output error levels for predictive assessments of compound-protein interactions.

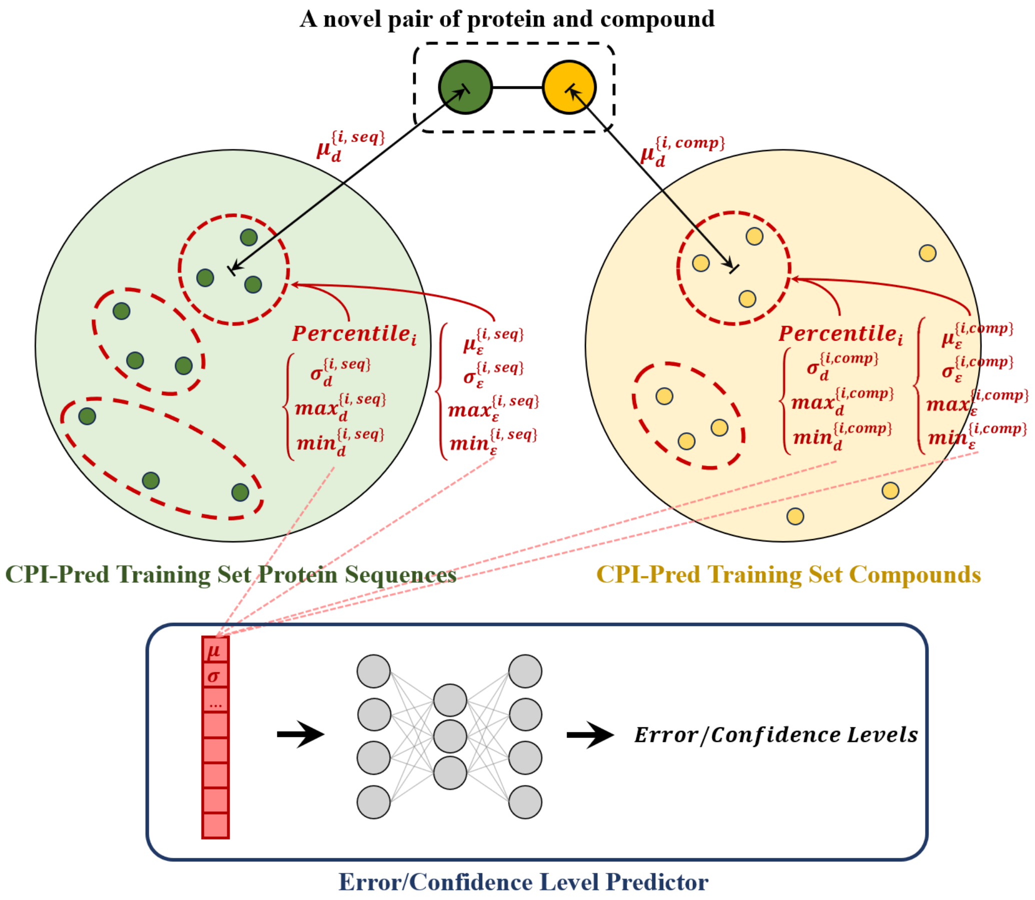


Figure S4: Schematic representation of the error level predictor for compound-protein interaction predictions (CPI-Pred). The illustration details the methodology for calculating Euclidean distances between a novel pair of protein and compound, and all corresponding pairs in the CPI-Pred training set. Two separate distribution analyses are conducted: one for the protein sequences (green) and one for the compounds (yellow). Within each set, distances are calculated and then grouped into 100 percentile ranks, with key statistics—mean ($\mu$), standard deviation ($\sigma$), maximum ($max$), and minimum ($min$)—computed for each percentile group. These statistics characterize the distribution of distances and serve as inputs to a neural network that determines the prediction error levels.

#### Error Level Predictor Performance

Figure ?? presents confusion matrices illustrating the performance of our error prediction model across six different Enzyme Commission (EC) classes. The model shows varying degrees of accuracy, with some classes like (i.e., Lyases (Class 4), Isomerases (Class 5), and Ligases (Class 6) demonstrating higher prediction accuracy for high errors. In contrast, predictions of classes such as Oxidoreductases (Class 1), Transferases (Class 2), and Hydrolases (Class 3) are relatively more balanced across both error levels. This is attributed to the greater amount of data available in the training set for EC numbers 1, 2, and 3 compared to EC numbers 4, 5, and 6.

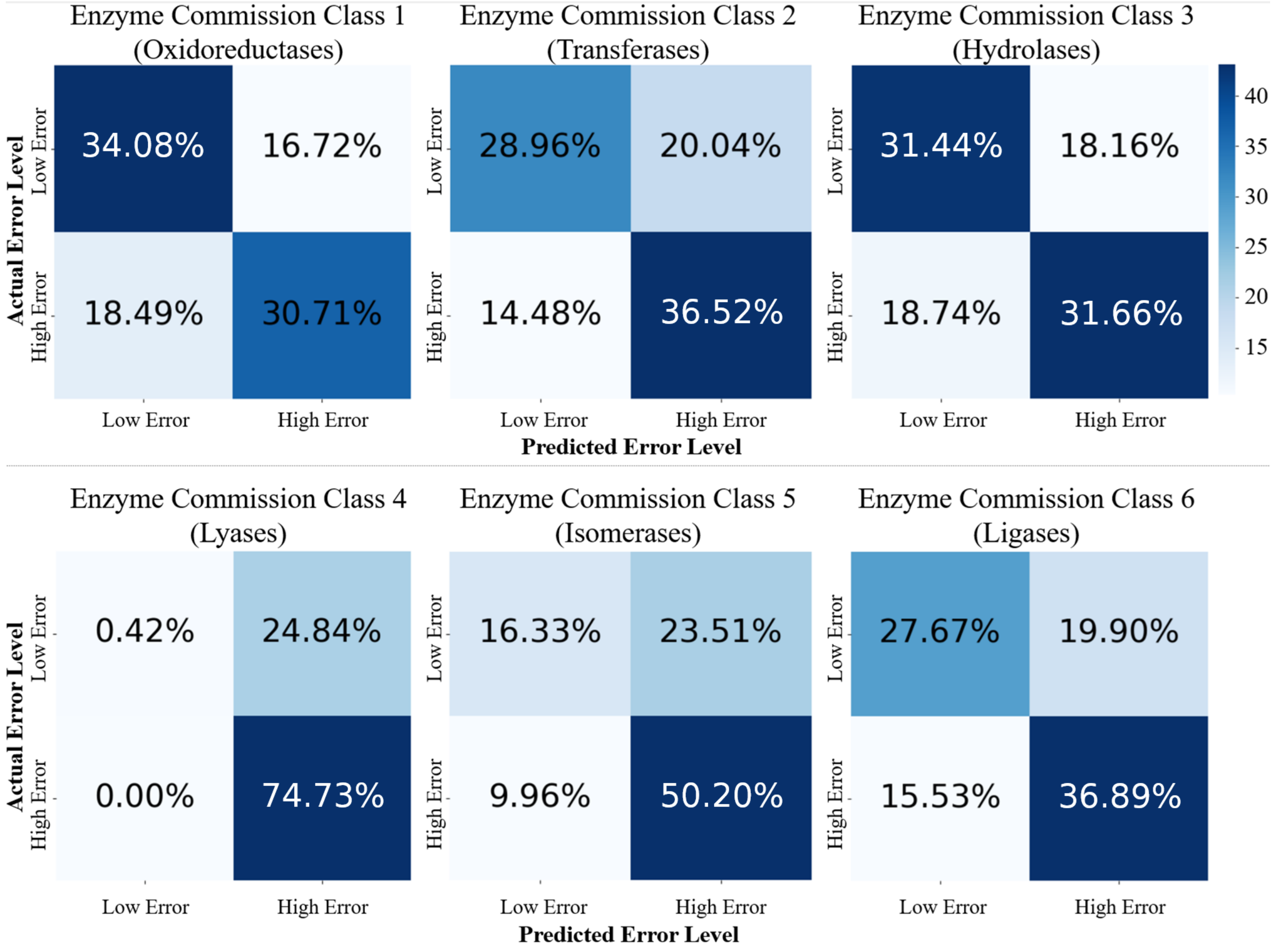


Figure S5: Error prediction performance across different enzyme commission classes

Figure S6 presents confusion matrices showing the performance of our error prediction model across three different data splitting tasks: random split, protein design, and compound design. The Random Split strategy achieves the highest F1 score of 0.779, compared to 0.374 for sequence design and 0.706 for compound design. This suggests that the model benefit from the more evenly distributed data in the random split.

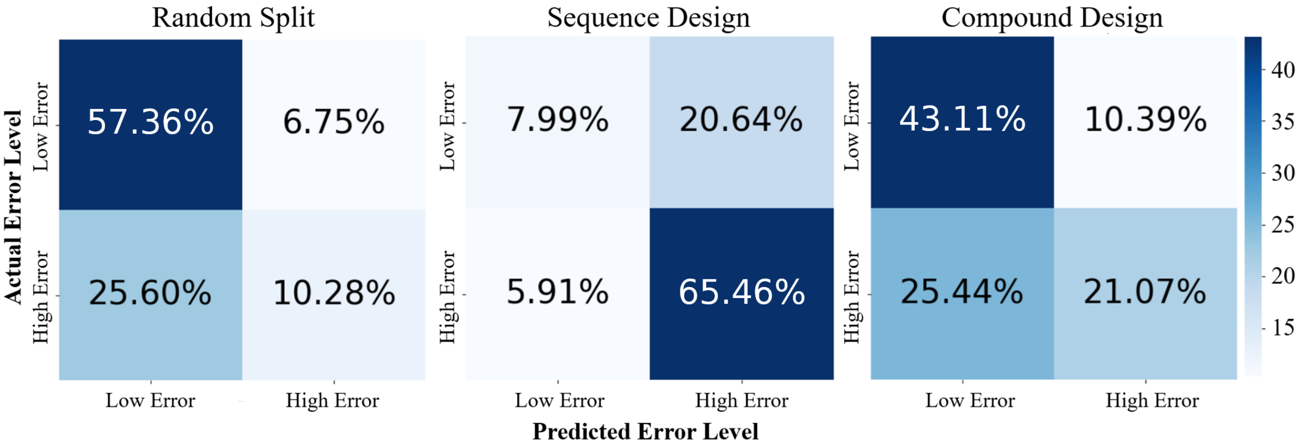


Figure S6: Error prediction performance across different data splitting tasks

### Performance of Different Sequence Embedding Methods

We evaluated four protein language models (ProtBERT[10], ESM-2[11], CARP[12], and Ankh[13]) and different sequence embedding pooling methods for sequence-to-function learning on two experimental datasets, PafA and avGFP. The enhanced performance of models utilizing convolutional or self-attention pooling signifies their promising utility in CPI prediction tasks. Detailed results are presented in Tables S3 and S4 as well as Figures S7 and S8.

Table S2: Comparisons of using different sequence embeddings and different embeddings pooling in downstream predictions with K-nearest neighbors being the baseline model. Models are trained and tested on *PafA* dataset containing kcat values for 1000 single point mutations. $R$ denotes Pearson's correlation and the best performing result is highlighted in bold.

| Sequence Embeddings | Training Database | Embeddings Dimension | Year Published |
|---|---|---|---|
| TAPE | Pfam | 768 | 2019 |
| ESM-1b | UniRef50 | 1280 | 2021 |
| ESM-1v | UniRef90 | 1280 | 2021 |
| ProtBERT | BFD100, UniRef100 | 1024 | 2021 |
| ESM-2 | UniRef50 | 5120 / 2560 / 1280 | 2022 |
| CARP | UniRef50 | 1280 | 2023 |
| Ankh | | 1536 | 2023 |

[a]Results being reported show average performances (and standard deviations) of models trained with five random seeds.

Table S3: Comparisons of using different language models and different embeddings pooling in downstream predictions (with K-nearest neighbors model being the baseline). Models are trained and tested on a *PafA* dataset containing experimental $k_{cat}$ values for 1034 point mutations (modifying or deleting one single amino acid in each sequence variants). "$r$" denotes Pearson's correlation coefficient and "$\rho$" denotes Spearman's rank correlation coefficient.The best performing result is highlighted in bold.

| Pooling Method Performance | Language Model | Self-Attention Pooling | Convolutional Pooling | LSTM-VAE Pooling | KNN (Baseline) |
|---|---|---|---|---|---|
| Pearson's $r$ | ESM-2 | **0.619±0.047** | 0.594±0.072 | 0.576±0.054 | 0.267±0.126 |
| | ESM-1v | 0.606±0.049 | 0.612±0.061 | 0.569±0.043 | |
| | ESM-1b | 0.616±0.046 | 0.608±0.041 | 0.571±0.060 | |
| | ProtBERT | 0.574±0.090 | 0.542±0.102 | 0.505±0.087 | |
| | Ankh | 0.560±0.051 | 0.545±0.044 | 0.506±0.054 | |
| | CARP | 0.421±0.070 | 0.460±0.048 | 0.435±0.041 | |
| Spearman's $\rho$ | ESM-2 | **0.641±0.039** | 0.600±0.059 | 0.583±0.048 | 0.234±0.143 |
| | ESM-1v | 0.610±0.055 | 0.636±0.057 | 0.577±0.040 | |
| | ESM-1b | 0.613±0.042 | 0.606±0.036 | 0.560±0.072 | |
| | ProtBERT | 0.588±0.073 | 0.569±0.092 | 0.509±0.092 | |
| | Ankh | 0.577±0.061 | 0.562±0.049 | 0.517±0.063 | |
| | CARP | 0.423±0.069 | 0.451±0.050 | 0.429±0.060 | |

Table S4: Comparisons of using different language models and different embeddings pooling in downstream predictions (with eUniRep in Low-N prediction model[14] serving as a baseline). Models are trained and tested on *avGFP* dataset. "$r$" denotes Pearson's correlation coefficient and "$\rho$" denotes Spearman's rank correlation coefficient. The best performing result is highlighted in bold.

| Metrics | Model | Self-Attention | Conv-1D pooling | LSTM-VAE | Low-N/eUniRep |
|---|---|---|---|---|---|
| Pearson's $r$ | ESM-2 | 0.926±0.001 | 0.962±0.001 | 0.863±0.006 | 0.871±0.002 |
| | ProtBERT | 0.910±0.004 | 0.950±0.004 | 0.857±0.007 | |
| | Ankh | 0.925±0.005 | **0.964±0.003** | 0.812±0.011 | |
| | CARP | 0.902±0.003 | 0.890±0.003 | 0.795±0.020 | |
| Spearman's $\rho$ | ESM-2 | 0.825±0.004 | **0.865±0.003** | 0.747±0.004 | 0.798±0.002 |
| | ProtBERT | 0.818±0.006 | 0.852±0.005 | 0.721±0.015 | |
| | Ankh | 0.820±0.004 | 0.858±0.005 | 0.704±0.009 | |
| | CARP | 0.815±0.003 | 0.796±0.002 | 0.699±0.018 | |

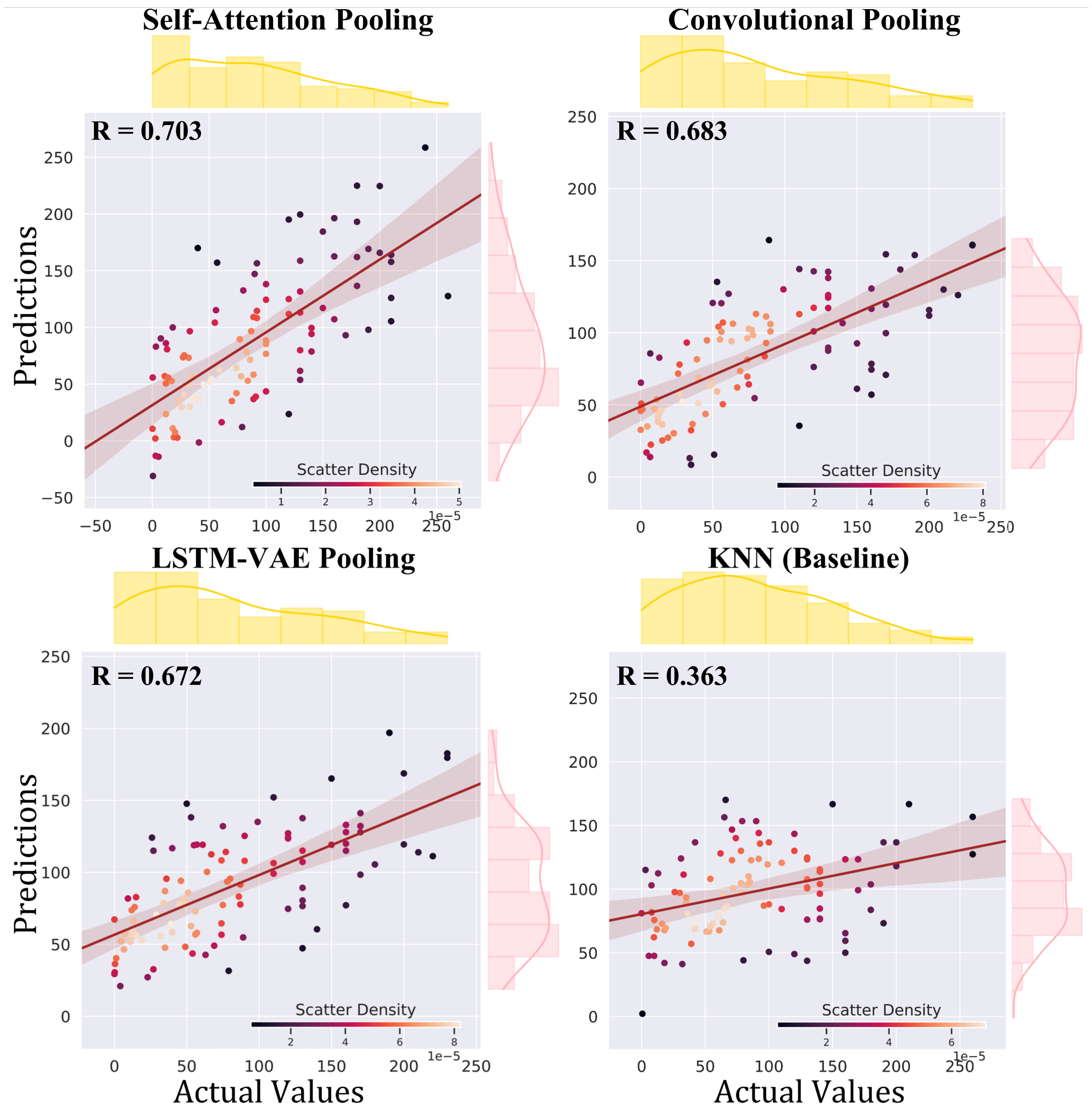


Figure S7: Visual comparison of different sequence embedding pooling methods including self-attention pooling, convolutional pooling, LSTM-VAE pooling on *PafA* dataset, with K-nearest neighbors (KNN) serving as a baseline. All methods use the same protein language model, ESM-2, for generating embeddings. The scatter plots shows predicted values versus actual values, providing a direct visual representation of the prediction capabilities of each method. Self-attention pooling yields the best predictive performance among all the evaluated pooling strategies.

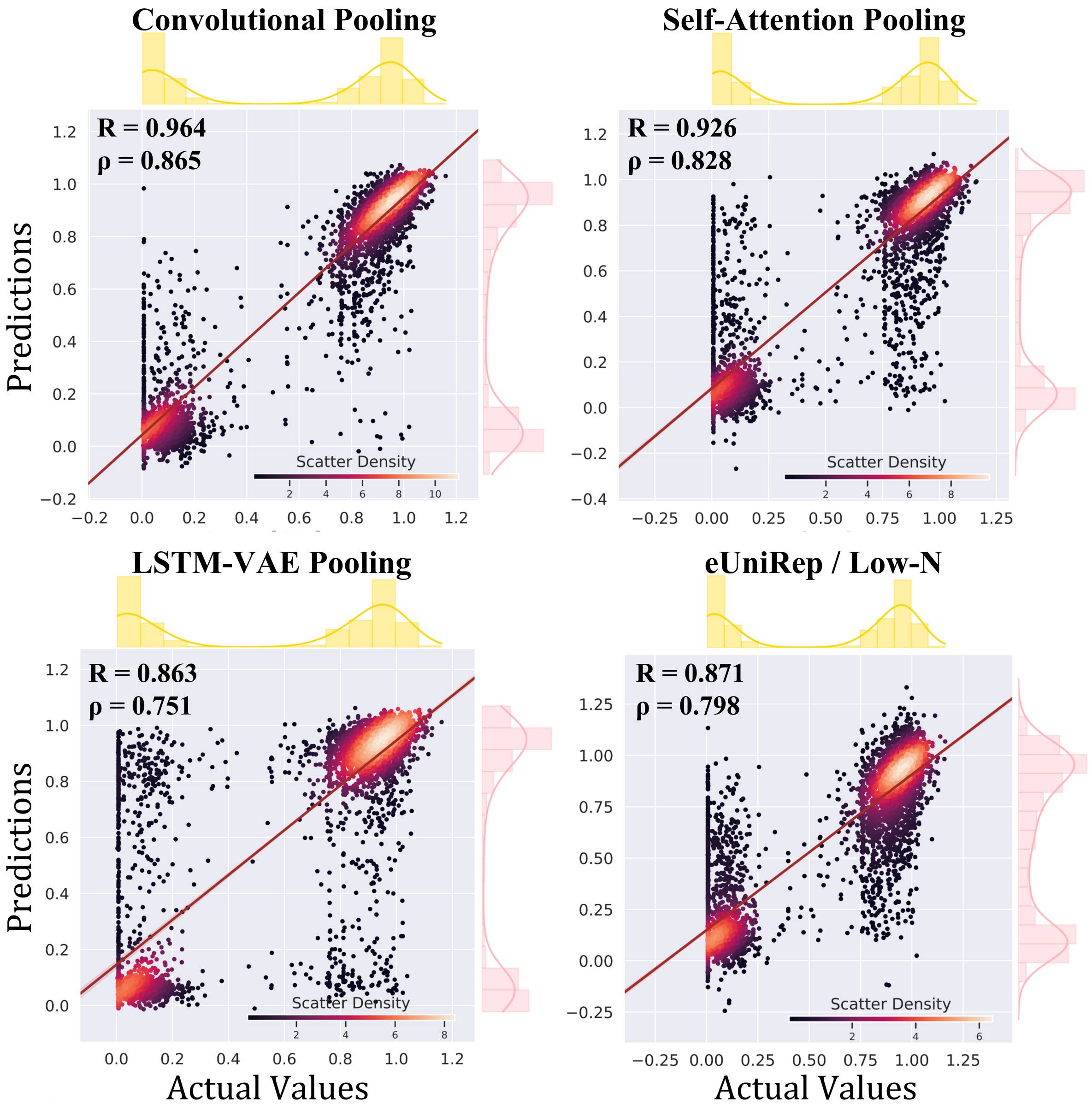


Figure S8: Visual comparison of different sequence embedding pooling methods including self-attention pooling, convolutional pooling, LSTM-VAE pooling on *avGFP* dataset, with eUniRep (in Low-N prediction model[14]) serving as a baseline. All methods use the same protein language model, ESM-2, for generating embeddings. The scatter plots shows predicted values versus actual values, providing a direct visual representation of the prediction capabilities of each method. Convolutional pooling yields the best predictive performance among all the evaluated pooling strategies and outperforms the eUniRep model.

### Performance of Different Compound Encoders in CPI Prediction

The improved compound encoding MPNN (with novel atom-level featurization) used in CPI-Pred demonstrates strong predictive performance, as evidenced in Table S5. It consistently outperforms other molecule encoding methods such as d-MPNN[15], ECFP6 count encoding[16], and Morgan fingerprint bit vector representation on $k_{cat}$ and $K_I$ core datasets. Statistical significance is assessed using a two-tailed Student's t-test at the 95% confidence level (* denotes $p < 0.05$, ** denotes $p < 0.01$), highlighting the robustness of the MPNN encoding's predictive capabilities.

Table S5: Comparison over different molecule encodings. Statistical significance is assessed using a two-tailed Student's t-test at the 95% confidence level (* denotes $p < 0.05$, ** denotes $p < 0.01$)

| Molecule Encodings | Dataset #Data | $k_{cat}$ (Core) 11,834 | | $K_I$ (Core) 4,341 | |
|---|---|---|---|---|---|
| **MPNN** w/ substructures count encodings | Pearson's $r$ | **0.781 ± 0.007** | ** | **0.751 ± 0.023** | ** |
| | Spearman's $\rho$ | 0.778 ± 0.013 | ** | 0.749 ± 0.038 | ** |
| | RMSE | 0.915 ± 0.010 | ** | 1.042 ± 0.021 | ** |
| d-MPNN | Pearson's $r$ | 0.762 ± 0.012 | * | 0.705 ± 0.032 | |
| | Spearman's $\rho$ | 0.757 ± 0.019 | | 0.697 ± 0.040 | |
| | RMSE | 1.001 ± 0.036 | | 1.198 ± 0.053 | |
| ECFP6 Count Encodings | Pearson's $r$ | 0.770 ± 0.005 | ** | 0.723 ± 0.027 | * |
| | Spearman's $\rho$ | 0.766 ± 0.011 | * | 0.733 ± 0.028 | * |
| | RMSE | 0.917 ± 0.013 | ** | 1.086 ± 0.025 | |
| Morgan Fingerprint Bit Vector | Pearson's $r$ | 0.753 ± 0.017 | | 0.718 ± 0.016 | |
| | Spearman's $\rho$ | 0.743 ± 0.020 | | 0.724 ± 0.037 | |
| | RMSE | 1.007 ± 0.043 | | 1.144 ± 0.021 | |

### Performance of Sequence Encoding Study in CPI Prediction

In parallel to the compound encoding strategies, the efficacy of different sequence encoding blocks was evaluated in a study. These encodings were assessed across two core datasets $k_{cat}$ and $K_I$ to ascertain their performance in CPI predictions. The sequence encoding techniques tested include n-grams encoding, sequence embeddings flattened (no pooling), sequence embeddings with average pooling and embeddings with self-attention pooling. For all tests within this study, Morgan bit vector was employed for encoding the compounds to ensure consistency across all tests. This method was chosen over the superior-performing MPNN-learnt representation due to its widespread use in published work. As presented in Table S6, the performance of the self-attention pooling method matches the no pooling method quite closely while producing a model that it about 1% of the no pooling method model. This suggests that this pooling method conserves a significant amount of information obtained from the protein language models while efficiently processing and reducing the dimension of the information provided.

Table S6: Comparison over different sequence encodings. Statistical significance is assessed using a two-tailed Student's t-test at the 95% confidence level (* denotes $p < 0.05$, ** denotes $p < 0.01$). Morgan bit vector was used for compound encoding across all tests here, chosen for its commonality in published works.

| Sequence Encoding | Dataset #Data | $k_{cat}$ (Core) 11,834 | $K_I$ (Core) 4,341 | Model Size |
|---|---|---|---|---|
| Self-Attention | Pearson's $r$ | 0.766 ± 0.007 ** | 0.692 ± 0.017 ** | |
| Pooling | Spearman's $\rho$ | 0.758 ± 0.009 ** | 0.701 ± 0.013 * | 4.1 M |
| Embeddings | RMSE | 0.969 ± 0.014 ** | 1.125 ± 0.031 ** | |
| Average | Pearson's $r$ | 0.656 ± 0.009 * | 0.674 ± 0.037 | |
| Pooling | Spearman's $\rho$ | 0.648 ± 0.011 * | 0.677 ± 0.049 | 3.4 M |
| Embeddings | RMSE | 1.140 ± 0.016 | 1.203 ± 0.069 * | |
| Flattened | Pearson's $r$ | 0.759 ± 0.012 ** | 0.623 ± 0.030 | |
| Embeddings | Spearman's $\rho$ | 0.753 ± 0.009 ** | 0.621 ± 0.031 | 335 M |
| (No Pooling) | RMSE | 0.967 ± 0.028 ** | 1.268 ± 0.088 | |
| N-gram | Pearson's $r$ | 0.588 ± 0.010 | 0.665 ± 0.022 | |
| Encoding | Spearman's $\rho$ | 0.586 ± 0.013 | 0.662 ± 0.031 | 0.3 M |
| | RMSE | 1.205 ± 0.028 | 1.291 ± 0.184 | |

### Bench-marking CPI-Pred

The tables presented in this section provide details of CPI-Pred's predictive performance against several state-of-the-art models across multiple tasks and datasets.

Table S7 extends the comparative performance analysis to five distinct enzyme activity datasets, namely phosphatase, halogenase, kinase, aminotransferase, and esterase. Here, the performance of CPI-Pred is contrasted with ESA-Pred[16], KNN, and DLkcat. CPI-Pred's predictions generally surpass all the latest models such as ESA-Pred and DLkcat.

Training CPI-Pred on a single kinetic parameter dataset demands between 24 and 96 hours of processing time on a single A100 GPU.

Table S7: Prediction performance (Pearson's correlation coefficients) of CPI-Pred on different tasks of five enzyme activity datasets, benchmarked against KNN and DLkcat. The shown results correspond to averages and standard deviations across models trained using 5-fold cross-validation.

| Tasks | Method | Phosphatase | Halogenase | Kinase | Aminotransferase | Esterase |
|---|---|---|---|---|---|---|
| Simple Task | CPI-Pred | **0.825**±0.012 | 0.886±0.049 | **0.858**±0.035 | **0.865**±0.044 | **0.853**±0.019 |
| | ESA-Pred | 0.816±0.020 | **0.892**±0.056 | 0.845±0.041 | 0.838±0.039 | 0.828±0.020 |
| | KNN | 0.563±0.031 | 0.535±0.022 | 0.537±0.060 | 0.604±0.052 | 0.765±0.032 |
| | DLkcat | 0.623±0.078 | 0.680±0.085 | 0.628±0.019 | 0.520±0.150 | 0.817±0.028 |
| Compound Design | CPI-Pred | 0.689±0.049 | 0.552±0.042 | 0.390±0.040 | 0.472±0.072 | 0.776±0.022 |
| | ESA-Pred | 0.681±0.036 | 0.545±0.070 | 0.335±0.067 | 0.470±0.081 | 0.751±0.019 |
| | KNN | 0.543±0.037 | 0.420±0.048 | 0.262±0.028 | 0.541±0.067 | 0.632±0.027 |
| | DLkcat | 0.569±0.072 | 0.435±0.072 | 0.366±0.036 | 0.520±0.073 | 0.673±0.025 |
| Protein Design | CPI-Pred | 0.466±0.092 | 0.670±0.042 | 0.760±0.044 | 0.798±0.062 | 0.359±0.068 |
| | ESA-Pred | 0.465±0.089 | 0.673±0.043 | 0.735±0.051 | 0.790±0.050 | 0.351±0.070 |
| | KNN | 0.269±0.061 | 0.512±0.069 | 0.619±0.033 | 0.502±0.080 | 0.225±0.042 |
| | DLkcat | 0.351±0.076 | 0.581±0.078 | 0.545±0.050 | 0.617±0.042 | 0.296±0.037 |

### Bench-marking via Comparison with ESP

In this comparative evaluation, we contrast our CPI-Pred model with the recently published ESP (Enzyme Substrate Prediction) model[17], which was released after the submission of our manuscript. ESP is a classification model that predicts the likelihood of a compound being a substrate for a given enzyme. Despite some methodological similarities with CPI-Pred, ESP does not incorporate advanced sequence embedding pooling, enhanced compound encoding, or our cross-attention based interaction prediction. The advancements brought by these techniques used by CPI-Pred are already illustrated in Sections D and E (e.g., the comparison between the compound encoder used by CPI-Pred and the "d-MPNN" used by ESP is shown in TableS5).

We compare the models using our extensive pangenomic datasets (including $k_{cat}$, $K_I$, $K_M$). ESP was trained on an expanded $K_M$ dataset, supplemented with similar compounds as negative data points, but only offers this $K_M$ dataset for comparison. Conversely, our pangenomic $K_M$ dataset, built based on KinMod, includes more phylogenetically inferred evidence and is larger than ESP's expanded $K_M$ dataset ($\sim$ 88,000 against $\sim$ 69,500). For a broader evaluation and to assess model versatility, we also considered the pangenomic $k_{cat}$ and $K_I$ datasets.

As shown in Figure S9, both models perform well, but CPI-Pred slightly outperforms ESP for enzyme-substrate classification, as indicated by higher ROC-AUC and AU-PRC metrics. This comparative evaluation underscores the methodological advantages of CPI-Pred, including the integration of advanced encoding and pooling techniques, and the use of larger, more comprehensive datasets. These features contribute to CPI-Pred's improved performance in predictive protein engineering tasks.

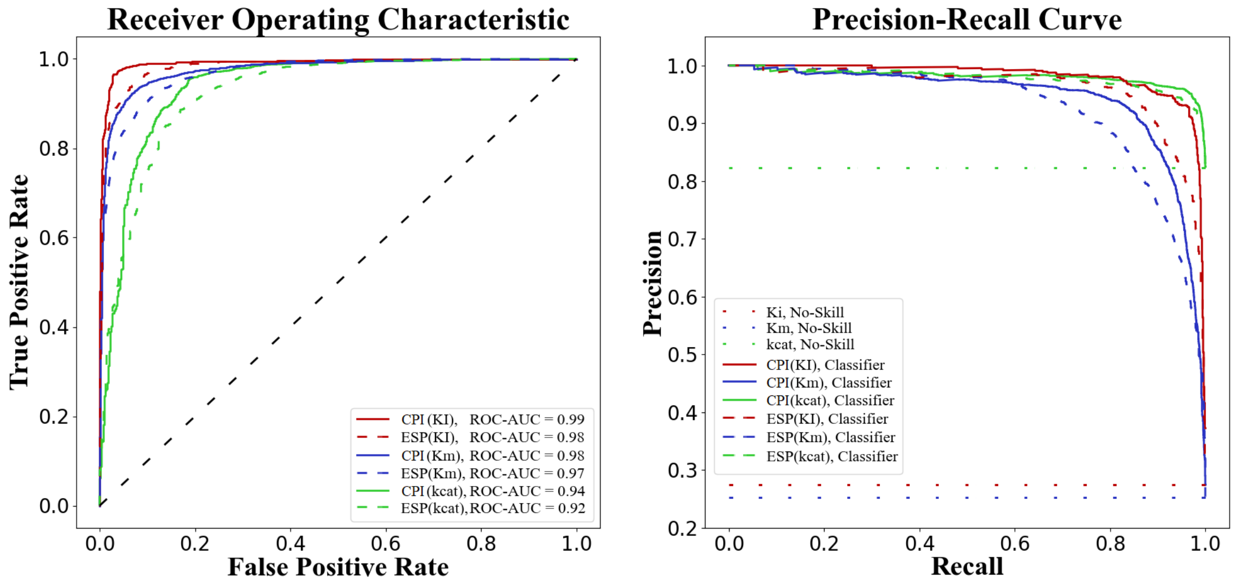


Figure S9: Visual comparison between the CPI-Pred and ESP models on the prediction of kinetic parameters $K_I$, $k_{cat}$, and $K_M$. The left panel presents the Receiver Operating Characteristic (ROC) curves, with the Area Under Curve (AUC) scores delineated in the legend. The right panel depicts the Precision-Recall (PRC) curves. In these plots, solid lines represent the performance of CPI-Pred, dashed lines indicate ESP, and dotted lines serve as a reference for a no-skill classifier. The kinetic parameters are color-coded: red for $K_I$, green for $k_{cat}$, and blue for $K_M$.

(10) Elnaggar, A.; Heinzinger, M.; Dallago, C.; Rehawi, G.; Yu, W.; Jones, L.; Gibbs, T.; Feher, T.; Angerer, C.; Steinegger, M.; Bhowmik, D.; Rost, B. ProtTrans: Towards Cracking the Language of Lifes Code Through Self-Supervised Deep Learning and High

Performance Computing. *IEEE Transactions on Pattern Analysis and Machine Intelligence* **2021**, *14*, 1–16.

(11) Lin, Z.; Akin, H.; Rao, R.; Hie, B.; Zhu, Z.; Lu, W.; dos Santos Costa, A.; Fazel-Zarandi, M.; Sercu, T.; Candido, S.; Rives, A. Language models of protein sequences at the scale of evolution enable accurate structure prediction. *bioRxiv* **2022**,

(12) Yang, K. K.; Fusi, N.; Lu, A. X. Convolutions are competitive with transformers for protein sequence pretraining. Pages: 2022.05.19.492714 Section: New Results.

(13) Elnaggar, A.; Essam, H.; Salah-Eldin, W.; Moustafa, W.; Elkerdawy, M.; Rochereau, C.; Rost, B. Ankh: Optimized Protein Language Model Unlocks General-Purpose Modelling. Issue: arXiv:2301.06568.

(14) Biswas, S.; Khimulya, G.; Alley, E. C.; Esvelt, K. M.; Church, G. M. Low-N protein engineering with data-efficient deep learning. *Nature Methods* **2021**, *18*, 389–396.

(15) Yang, K.; Swanson, K.; Jin, W.; Coley, C.; Eiden, P.; Gao, H.; Guzman-Perez, A.; Hopper, T.; Kelley, B.; Mathea, M.; Palmer, A.; Settels, V.; Jaakkola, T.; Jensen, K.; Barzilay, R. Analyzing Learned Molecular Representations for Property Prediction. *Journal of Chemical Information and Modeling* **2019**, *59*, 3370–3388, PMID: 31361484.

(16) Xu, Z.; Wu, J.; Song, Y. S.; Mahadevan, R. Enzyme Activity Prediction of Sequence Variants on Novel Substrates using Improved Substrate Encodings and Convolutional Pooling. Proceedings of the 16th Machine Learning in Computational Biology meeting. 2022; pp 78–87.

(17) Kroll, A.; Ranjan, S.; Engqvist, M. K. M.; Lercher, M. J. A general model to predict small molecule substrates of enzymes based on machine and deep learning. *Nature Communications* **2023**, *14*, 2787.
